# An intron-based timer for circadian rhythms

**DOI:** 10.1101/2025.10.27.684876

**Authors:** Ye Yuan, Amanda Linskens, Rafael De Gouvea, Haokai Liu, Yangbo Xiao, Smriti Suresh, Irene Hu, Swathi Yadlapalli

## Abstract

**Abstract:** Circadian clocks regulate daily rhythms in all eukaryotes through ∼24-hour transcriptional-translational feedback loops driven by clock proteins. However, the molecular mechanisms that set the 24-hour period remain poorly defined. Here, using single-molecule imaging and nascent RNA sequencing, we uncover an unexpected RNA-based molecular timer: a single intron in the *Drosophila timeless* (*tim*) gene that regulates circadian period length by controlling mRNA localization. Strikingly, we find that ∼50% of *tim* mRNAs are localized to the nucleus due to inefficient post-transcriptional splicing of a single intron (which we named *intron P*), in contrast to other core clock transcripts that localize predominantly to the cytoplasm. CRISPR-mediated removal of intron P abolishes nuclear retention of *tim* transcripts, leading to accelerated TIM protein accumulation and a shortened ∼22-hour period with reduced rhythmic robustness. Remarkably, insertion of intron P alone into heterologous reporters is sufficient to promote nuclear retention in both *Drosophila* and human cells, acting as a conserved checkpoint that withholds transcripts in the nucleus until splicing is complete. Finally, we identify three RNA-binding proteins, two repressors (Hrb27C and Squid) and one activator (Qkr58E-2, a Sam68 homolog), that modulate intron P splicing in a rheostat-like manner. Together, these findings establish *tim* intron P as the first intron-based molecular timer in circadian clocks and reveal splicing kinetics as a critical regulatory layer in temporal gene expression programs, with broad implications for other processes such as development and immunity.

## Introduction

The ability of circadian clocks to generate ∼24-hour rhythms underpins daily physiology and behavior in all eukaryotes. At their core, circadian clocks are built on transcriptional– translational feedback loops (TTFLs), in which clock proteins repress their own transcription and thereby drive self-sustained oscillations (Extended Data Fig. 1A) ^1, 2^. A central, long-standing question in chronobiology is what molecular mechanisms set the ∼24-hour period length. Models to explain circadian timing have traditionally relied on protein-based mechanisms, such as regulated degradation of PERIOD ^3, 4^ and TIMELESS ^5^ or the slow enzymatic cycles of KaiC ^6^ and RUVBL2 ^7^, as the key determinants of the 24-hour period. However, whether RNA-level regulation can control 24-hour circadian period remains unknown.

While alternative splicing is well known to diversify transcript and protein isoforms, much less is understood about how the kinetics of intron removal and nuclear export shape gene expression timing ^8^. RNA processing events—capping, splicing, and polyadenylation—are often assumed to occur rapidly and co-transcriptionally, ensuring efficient delivery of mature transcripts to the cytoplasm in most models of gene expression ^9^. However, recent transcriptomic studies in human, mouse, and plant cells have revealed that many polyadenylated transcripts retain introns and remain in the nucleus ^10–12^. These findings raise the possibility that post-transcriptional splicing and nuclear retention could serve as regulatory checkpoints, but whether such mechanisms contribute to circadian timing has not been established.

To systematically investigate whether nuclear RNA processing can serve as a rate-limiting step in the circadian clock, we analyzed the splicing and localization dynamics of core clock mRNAs in *Drosophila* clock neurons (Extended Data Fig. 1B). Here we report that inefficient splicing of a single intron in the *Drosophila timeless* (*tim*) gene, which we call intron P, acts as a molecular timer that fine-tunes circadian period length. We demonstrate that intron P functions as a conserved nuclear export checkpoint, retaining transcripts in the nucleus until splicing is complete and thereby controlling the timing and levels of TIM protein accumulation, and ultimately, the pace of circadian oscillations. These findings reveal splicing kinetics as a critical rate-limiting step in circadian timing and highlight introns as potential dynamic regulators of temporal gene expression in diverse biological contexts.

### A single retained intron regulates nuclear localization of timeless mRNAs

To test whether nuclear RNA processing acts as a regulatory checkpoint in circadian timing, we first examined the subcellular localization of core clock mRNAs. We reasoned that if post-transcriptional splicing or nuclear export serves as a regulatory bottleneck, we would observe differential accumulation of these mRNAs in the nucleus compared to the cytoplasm. To visualize mRNAs with subcellular resolution, we adapted the Hybridization Chain Reaction-Fluorescence In Situ Hybridization (HCR-FISH) technique, which enables the detection of individual RNA molecules as diffraction-limited spots ^13^. For each of the core clock genes, we designed FISH probes targeting a common exon across all isoforms (see Methods). We uncovered a striking localization pattern for *timeless* (*tim*) mRNAs: Unlike other core clock mRNAs like *period* and *cryptochrome*, which are localized exclusively in the cytoplasm (Extended Data Fig. 1C-E), *tim* mRNAs were found in both the nucleus and the cytoplasm throughout the circadian cycle (Fig. 1A). We quantified the number of *tim* mRNA spots in large lateral ventral neurons (l-LNvs), a key clock neuron group that controls rhythmic behavior ^2^, and found that while their levels oscillate (Fig. 1B), the percentage of *tim* mRNAs that are localized to the nucleus remained steady at ∼50% throughout the circadian cycle (Fig. 1C). In *per^01^*null mutants, which lack a functional clock ^14^, *tim* mRNA oscillation is disrupted; however, the ∼50:50 nucleo-cytoplasmic distribution persisted (Extended Data Fig. 1F-H), indicating that the subcellular distribution of *tim* mRNAs is clock independent. To further validate these findings, we performed single molecule FISH (smFISH) experiments ^15^ using probes spanning the entire *tim* coding sequence (Extended Data Fig. 1I-K). Our experiments using smFISH and HCR-FISH techniques produced similar results regarding the total number and the nuclear localization of *tim* mRNAs.

**Fig. 1.**
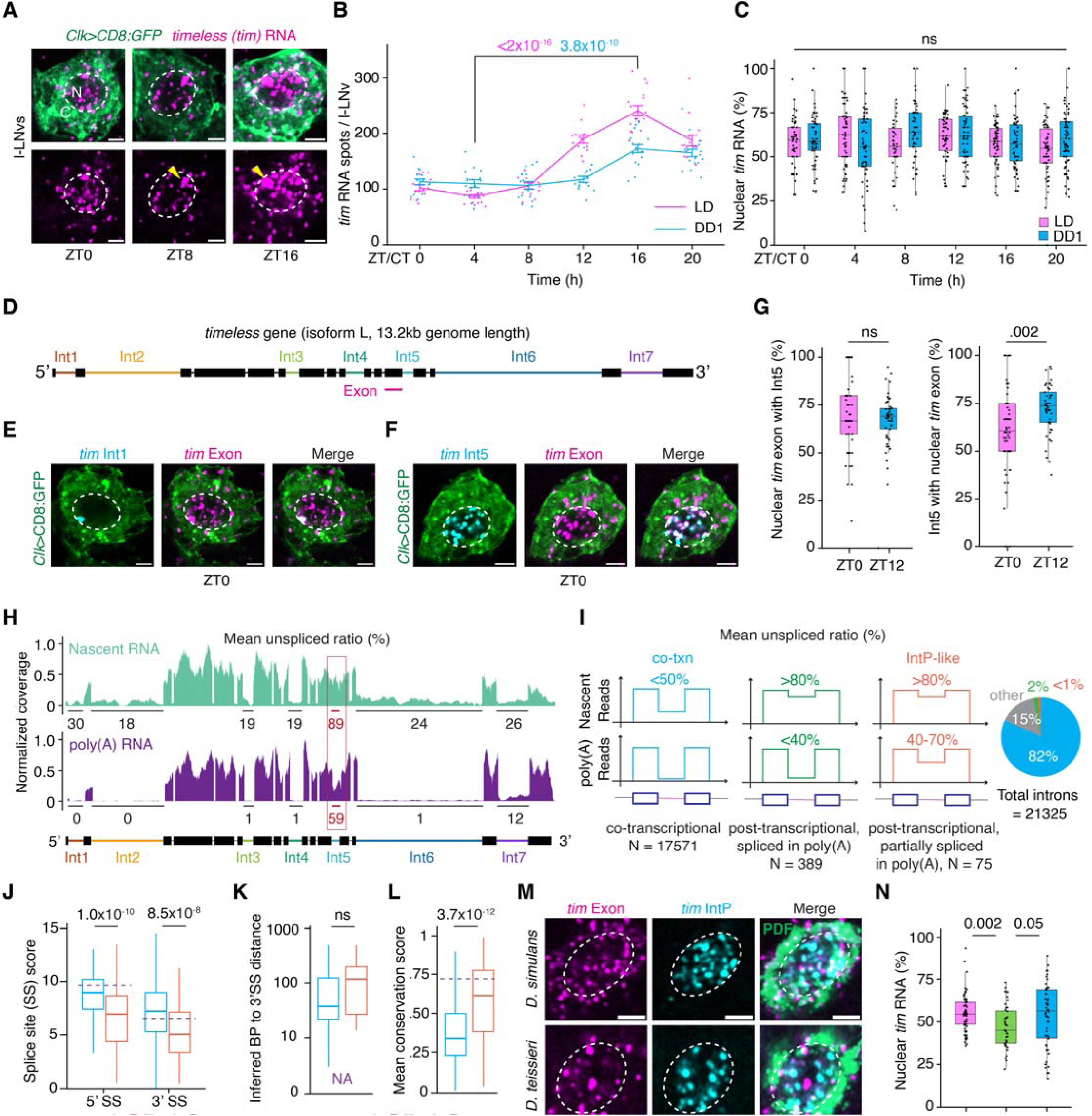
Intron P of *timeless* undergoes post-transcriptional splicing. (**A**) Representative images of *timeless* (*tim*) RNA in l-LNv neurons at different time points (ZT0, ZT8, ZT16) across the circadian cycle. l-LNv cytoplasm is shown in green, *tim* RNA spots are shown in magenta, and nuclei are outlined in white. ZT0 represents the repression phase; ZT8 and ZT16 represent the activation phase (see Methods). ‘N’ denotes nucleus, ‘C’ denotes cytoplasm, and yellow arrowheads indicate transcription sites. (**B**) Quantification of *tim* RNA spot counts per l-LNv over the light-dark cycle (LD, pink trace) and the first day of constant darkness (DD1, blue trace). P-values for RNA levels at ZT4 versus ZT16 are shown in pink for LD and in blue for DD1. (**C**) Quantification of the percentage of nuclear *tim* RNA in l-LNvs over LD (pink) and DD1 (blue). (**D**) Schematic of the *tim* RNA (isoform L) with all intron probes and the exon probe used to detect mature *tim* RNA highlighted. (**E**) Representative image of *tim* intron 1 (cyan) and exon (magenta) FISH signals in a l-LNv. (**F**) Representative image of *tim* intron 5 (cyan) and exon (magenta) FISH signals in a l-LNv. (**G**) Percentage of nuclear *tim* exon spots colocalizing with intron-5 spots (left panel) and intron 5 spots colocalizing with nuclear *tim* exon spots (right panel) in l-LNvs during repression (ZT0, pink box plots) and activation (ZT12, blue box plots) phases. (**H**) Normalized read coverage of the *tim* locus from Nascent-seq (top) and poly(A)-seq (bottom). Splicing status of *tim* introns, measured as mean unspliced read ratios, is shown as percentages. The mean unspliced ratio for intron 5 (Intron P) is highlighted in red. (**I**) Schematic of genome-wide classification of intron splicing events as ‘co-transcriptional’ (co-txn, blue) or ‘post-transcriptional’ (green and orange). The mean unspliced ratio cutoffs used to define these groups are shown above the plots. Introns with mean unspliced ratios similar to *tim* Intron P are labeled ‘IntP-like’ (orange). A pie chart summarizing the classification results is shown on the right. (**J-L**) Analysis of 5’ and 3’ splice site strength (**J**), distance from inferred branch point to the 3’ splice site (**K**), and mean conservation score (**L**) for introns belonging to the ‘co-txn’ group (blue box plots) and the ‘IntP-like’ group (orange box plots). Intron P scores are highlighted as purple lines. NA, not applicable. (**M**) Representative images of *D. mel. tim* exon (magenta) and *D. mel. tim* Intron P (cyan) FISH signals in l-LNvs from *Drosophila simulans* (*D. sim*) and *Drosophila teissieri* (*D. tei*) flies. l-LNv cytoplasm is marked by PDF antibody (green). (**N**) Quantification of the percentage of nuclear *tim* RNA in l-LNvs in all three *Drosophila* species. In **B**, data are presented as line plots showing the mean ± standard error of the mean (SEM). Statistical analysis was performed using a two-sided Student’s t-test. In **C**, **G**, **J**, **K**, **L**, and **N**, data are shown as box plots displaying the median and quantiles. Statistical analysis was performed using two-sided Mann-Whitney U tests. ns; not significant. In all plots, each dot represents data from l-LNvs of a single *Drosophila* hemi-brain. Refer to Supplementary Table S10 for all statistical comparisons. l-LNv nuclei are circled in white. Scale bars-2µm.

In our HCR-FISH experiments using the *tim* exon probe, we consistently detected a single, highly concentrated, non-diffraction-limited spot (yellow arrowhead in Fig. 1A), especially during the activation phase (ZT8, ZT16), indicating that multiple nascent RNAs are being synthesized simultaneously at the gene’s transcription site ^16^ (Fig. 1A). In addition, we observed several distinct spots distributed throughout the nucleus. We hypothesized that these nuclear *tim* mRNA spots might correspond to partially spliced mRNAs retaining specific introns. To test this hypothesis, we designed FISH probes targeting each of the *tim* introns (excluding those <100 bp) and performed dual-color HCR-FISH experiments with both intron and exon probes. We found that the majority of the introns (1, 2, 3, 4, and 6) predominantly appeared as a single nuclear spot that colocalized with the brightest exon signal, corresponding to the transcription site (Fig. 1D, E; Extended Data Fig. 2A-B). This finding indicates that these introns are being spliced out during transcription, consistent with co-transcriptional splicing ^16^. In striking contrast, intron 5—henceforth referred to as ‘Intron P’ (P for ‘post-transcriptional’)— exhibited a disperse nuclear pattern, suggesting it is not spliced out co-transcriptionally (Fig. 1F). We observed a similar localization pattern for *tim* exon and Intron P in other clock neuron groups (Extended Data Fig. 2C). We observed that a majority of the nuclear *tim* exon spots (>70%) co-localized with Intron P spots in l-LNvs, while the remaining (∼30%) likely represent fully spliced transcripts poised for export (Fig. 1G, left panel; Extended Data Fig. 2D). Similarly, ∼75% of the Intron P spots co-localized with the nuclear exon spots, with the remainder likely corresponding to excised introns awaiting degradation (Fig. 1G, right panel; Extended Data Fig. 2D). Importantly, Intron P-containing transcripts were never detected in the cytoplasm, implying that Intron P is spliced out before export. To further corroborate our imaging-based observations, we performed long-read sequencing of polyadenylated RNAs from *Drosophila* brains, which confirmed that ∼50% of mature *tim* transcripts retain Intron P (Extended Data Fig. 2E).

To understand how Intron P might contribute to the ∼50:50 nuclear to cytoplasmic distribution of *tim* mRNAs, we considered three possible models: (1) Intron P is always retained during transcription and is subsequently spliced post-transcriptionally before export; (2) Intron P is spliced co-transcriptionally in some transcripts, which are exported, while it remains unspliced in others, causing them to remain nuclear; or (3) Intron-P-containing transcripts, which harbor a premature stop codon, are exported and degraded by the Nonsense-Mediated Decay (NMD) pathway ^17^. To distinguish between these possibilities, we analyzed a published nascent RNA-seq dataset from *Drosophila* brains ^18^. We found that ∼90% of newly synthesized *tim* mRNAs include Intron P— far higher than the < 20% inclusion seen for other introns—indicating that most nascent *tim* transcripts retain Intron P (Fig. 1H, top). Intron P levels decrease to ∼50% in the mature, polyadenylated fraction, suggesting that this intron is spliced out post-transcriptionally in some of the transcripts (Fig. 1H, bottom), consistent with model 1.

To test the possibility whether Intron-P-containing transcripts could be exported and subsequently degraded by the NMD pathway (model 3), we knocked down three core NMD factors, Upf1, Upf2, and Upf3, in clock neurons using functionally validated RNAi fly lines ^19^. We found that clock neuron-specific knockdown of two of these core factors, Upf1 and Upf3, led to behavioral arrhythmicity (Extended Data Fig. 2F-G), suggesting that the NMD pathway plays an important role in circadian regulation. If NMD were responsible for degrading cytoplasmic *tim* transcripts containing Intron P, then inhibiting NMD would be expected to result in their cytoplasmic accumulation. Interestingly, previous transcriptomic studies detected Intron-P-retaining *tim* transcripts (termed “*tim-tiny*” ^20^ or “intron M” ^21^), but could not detect the corresponding protein, leading them to suggest that these transcripts might be degraded *via* the NMD pathway. Contrary to these expectations, we found no evidence of cytoplasmic accumulation of intron-P-retaining *tim* mRNAs upon knockdown of NMD pathway components (Extended Data Fig. 2H-J), indicating that NMD pathway does not regulate the nucleo-cytoplasmic distribution of *tim* transcripts. Together, these results unambiguously support model 1: Intron P is retained in nearly all nascent *tim* transcripts, is spliced post transcriptionally, and only fully spliced transcripts are exported.

Having established that *tim* Intron P is spliced post-transcriptionally, we next asked whether environmental cues critical for clock entrainment, such as light-dark cycles or temperature, modulate its splicing efficiency. We found that light-dark cycle cycles had no effect on *tim* mRNA splicing, as the nuclear-to-cytoplasmic ratios of *tim* mRNA remained consistent across the circadian cycle (Fig. 1C). Interestingly, splicing of Intron P was slightly more efficient at lower temperatures, as indicated by increased cytoplasmic accumulation of *tim* transcripts at 18 °C compared to 29 °C (Extended Data Fig. 3). These findings are consistent with prior RNA-seq studies showing higher Intron P retention at elevated temperatures ^21^. This temperature-dependent effect is also consistent with the known stabilization of RNA secondary structures at lower temperatures, which can enhance the spliceosome’s ability to recognize and process splice sites ^22^.

Next, we investigated the broader relevance of this post-transcriptional splicing mechanism by analyzing the nascent RNA-seq data from Drosophila brains ^18^. While a majority (80%) of introns are spliced co transcriptionally (Fig. 1I, ‘co-txn’ group), we identified 75 additional “IntP like” introns—including one in the synaptic gene *unc13* ^23^—that exhibit high retention in nascent transcripts but only moderate retention in the poly(A) fraction, suggesting similar post-transcriptional splicing dynamics (Fig. 1I, ‘IntP-like’ group; Extended Data Figs. 4A-C, 5A-B; Supplementary Tables S1, S2). HCR-FISH experiments showed that *unc13* transcripts, like *tim*, localize to both the nucleus and the cytoplasm in clock neurons, with nuclear exon signals colocalizing with intronic signals (Extended Data Fig. 4D-E). Sequence analysis revealed that most IntP-like introns possess “weak” splicing signals, suboptimal 5′ donor or 3′ acceptor sites (Fig. 1J) and/or branch points positioned far from the 3′ splice site (Fig. 1K), yet they are more highly conserved across *Drosophila* species (Fig. 1L; Extended Data Fig. 5C). Intriguingly, *tim* Intron P contains strong donor and acceptor splice sites but lacks a consensus branch point sequence (Fig. 1J, 1K, purple line). To test whether Intron P’s retention is a conserved feature, we conducted RNA-FISH experiments in two distantly related *Drosophila* species: *D. simulans*, which diverged ∼5.4 million years ago, and *D. teissieri*, which diverged ∼12.8 million years ago. In both species, *tim* transcripts were distributed both in the nucleus and cytoplasm of clock neurons, with Intron P signals confined to the nucleus (Fig. 1M, 1N). Together, these findings demonstrate that *tim* transcripts are localized to both the nucleus and the cytoplasm, with Intron P-containing transcripts retained in the nucleus and only fully spliced transcripts exported to the cytoplasm.

### Deletion of Intron P leads to a shortened circadian cycle with a period of ∼22.5 hours

To investigate the role of Intron P in circadian rhythms, we used CRISPR/Cas9 to excise it from the endogenous *tim* gene (*tim^intP^*^Δ^; Fig. 2A). We confirmed the intron knockout using FISH experiments, which showed no Intron P signals in the clock neurons of *tim^intP^*^Δ^ flies (Extended Data Fig. 6A). We found that while total *tim* mRNA levels in clock neurons, measured across peak and trough phases, remained comparable between wild-type and *tim^intP^*^Δ^ flies (Fig. 2B, 2C; Extended Data Fig. 6E), their subcellular distribution differed significantly (Fig. 2B). In wild-type flies, *tim* exon signals were evenly (∼50%) distributed between the nucleus and the cytoplasm (Fig. 2B, 2D). However, in *tim^intP^*^Δ^ flies the nuclear fraction dropped to less than 10% under both cycling light-dark and constant dark conditions (Fig. 2B, 2D; Extended Data Fig. 6C-E), as the remaining *tim* introns are spliced out co-transcriptionally (Extended Data Fig. 6B) and the fully spliced transcripts are rapidly exported to the cytoplasm. As a result, cytoplasmic *tim* mRNA levels in *tim^intP^*^Δ^ flies doubled compared to wild-type controls at all timepoints over the circadian cycle (Extended Data Fig. 6F-G). These results suggest that splicing of Intron P serves as a critical bottleneck for nuclear export, retaining *tim* transcripts in the nucleus until the intron is removed.

**Fig. 2.**
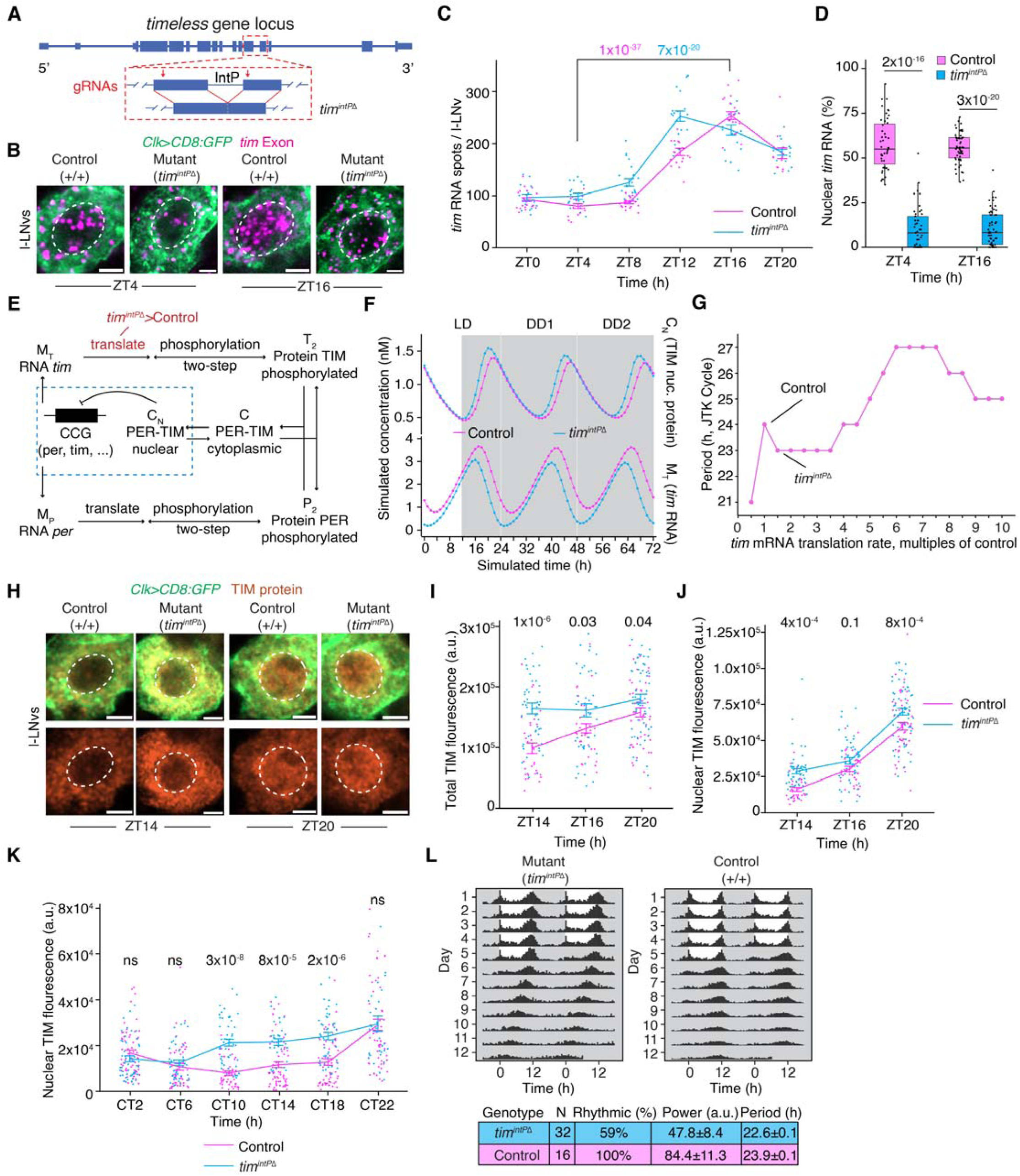
Deletion of Intron P eliminates nuclear retention of *tim* mRNAs and shortens the circadian cycle. (**A**) Schematic of the *tim* gene locus illustrating the CRISPR design used to delete Intron P (see Methods). (**B**) Representative images of *tim* exon spots in l-LNvs from control (Intron P intact) and *tim^IntP^*^Δ^ (Intron P deletion) flies at different time points across the light-dark cycle. l-LNv cytoplasm is shown in green, *tim* exon spots in magenta, and nuclei are outlined in white. (**C**) Quantification of *tim* RNA spot counts per l-LNv from control and *tim^IntP^*^Δ^ flies over the light-dark cycle (LD, pink trace) and the first day of constant darkness (DD1, blue trace). P-values for RNA levels at ZT4 *vs*. ZT16 are shown in pink (LD) and blue (DD1). (**D**) Quantification of the percentage of nuclear *tim* RNA in l-LNvs from control (pink box plots) and *tim^IntP^*^Δ^ (blue box plots) flies. (**E**) Schematic of a minimal model of the PER-TIM negative feedback. To account for elevated cytoplasmic *tim* mRNA in *tim^IntP^*^Δ^ flies, the *tim* mRNA translation rate was doubled in the *tim^IntP^*^Δ^ model (see Methods). (**F**) Simulated time series from the model showing nuclear TIM protein levels (top) and *tim* RNA levels (bottom) during the light-dark cycle (LD) and the first two days of constant darkness (DD1, DD2). Compared to the wild-type control, *tim^IntP^*^Δ^ exhibits earlier TIM protein accumulation and an advanced phase of *tim* RNA expression. (**G**) Simulated results from the model showing circadian clock period length as a function of *tim* RNA translation rate, analyzed using JTK_CYCLE. *tim^IntP^*^Δ^ shows a shorter period compared to the wild-type control. (**H**) Representative images of TIM protein immunostaining in l-LNvs from control and *tim^IntP^*^Δ^ flies at time points ZT14 and ZT20 (night) over the circadian cycle. l-LNv cytoplasm is shown in green, TIM protein in orange, and nuclei are outlined in white. (**I-J**) Quantification of the total TIM fluorescence intensity (**I**) and nuclear TIM fluorescence intensity (**J**) in l-LNvs from wild-type control and *tim^IntP^*^Δ^ flies at time points ZT14 and ZT20. (**K**) Quantification of nuclear TIM fluorescence intensity in l-LNvs from the wild-type control and *tim^IntP^*^Δ^ flies on the second day of constant darkness (DD2). (**L**) Averaged population locomotor-activity profiles of *tim^IntP^*^Δ^ flies (left panel) and control flies (right panel) in LD and DD conditions, with rest-activity patterns shown for two consecutive days in each row. Periodogram analysis under constant conditions is shown at the bottom. *tim^IntP^*^Δ^ flies exhibit a shortened circadian period of ∼22.5 hours and display less robust rhythms, as indicated by a lower percentage of rhythmic flies and reduced rhythm power, compared to control flies (pink). In **C**, **I**, **J**, **K,** data are presented as line plots showing the mean ± standard error of the mean (SEM). Statistical analysis was performed using a two-sided Student’s t-test. In **D**, data are shown as box plots displaying the median and quantiles. Statistical analysis was performed using two-sided Mann-Whitney U tests. ns, not significant. In all plots, each dot represents data from l-LNvs of a single *Drosophila* hemi-brain. Refer to Supplementary Table S10 for all statistical comparisons. l-LNv nuclei are circled in white. Scale bars-2µm.

To investigate how the elevated cytoplasmic *tim* mRNA levels in *tim^intP^*^Δ^ flies might affect the circadian clock, we employed in-silico simulations using an established model of the PER-TIM feedback loop ^24^ (see Methods, Fig. 2E; Extended Data Fig. 7A). Because the cytoplasmic *tim* mRNA sequence is identical in both wild-type and *tim^intP^*^Δ^ flies, it is reasonable to assume that the factors governing its stability and cytoplasmic degradation rates remain unchanged across conditions. Therefore, the elevated cytoplasmic mRNA levels in *tim^intP^*^Δ^ flies are expected to primarily enahnce TIM protein production. To model this, we doubled the translation rate of *tim* mRNAs in *tim^intP^*^Δ^ flies compared to the controls (Fig. 2E). Our simulations revealed that removing Intron P led to a) faster TIM protein accumulation (Fig. 2F, top panel), b) an earlier *tim* mRNA peak (Fig. 2F, bottom panel), and c) a 1-2 hour reduction in the circadian period (Fig. 2G). These simulations suggest that the clock runs faster in *tim^intP^*^Δ^ flies due to elevated cytoplasmic *tim* mRNA and increased translation. Notably, further increasing the translation rate in the model disrupted protein oscillations and reduced rhythmic robustness, indicating that excessive TIM production can destabilize the circadian clock’s regulatory dynamics (Extended Data Fig. 7B-E). Together, these results highlight the importance of tightly regulating TIM levels to maintain stable and robust circadian rhythms.

We experimentally tested the predictions from our model using *tim^intP^*^Δ^ flies. We found that TIM protein levels in clock neurons from *tim^intP^*^Δ^ flies rose earlier and peaked higher under both light-dark and constant darkness conditions (Fig. 2H, 2I; Extended Data Fig. 6H), consistent with increased translation driven by elevated cytoplasmic *tim* mRNA levels. We also observed higher nuclear TIM fluorescence in clock neurons from *tim^intP^*^Δ^ flies compared to controls. (Fig. 2J, 2K). Notably, TIM degradation timing, however, remained unaffected (Fig. 2K, Extended Data Fig. 6I-J), as excision of Intron P does not alter the TIM protein sequence or its interaction with CRY, which mediates light-dependent turnover ^25^. Additionally, clock neurons from *tim^intP^*^Δ^ flies exhibited an earlier peak in *tim* mRNA oscillation compared to wild-type controls (Fig. 2C), due to accelerated negative feedback from prematurely accumulated TIM protein. Finally, we observed that only ∼60% of *tim^intP^*^Δ^ flies maintained rhythmicity under constant darkness, with a shortened circadian period of ∼22.5 hours compared to ∼24 hours in controls (Fig. 2L), matching the model’s predicted 1–2-hour reduction. The reduced rhythmicity likely results from excessive TIM protein production, as variability in TIM levels among individual neurons may impair its ability to sustain a stable circadian cycle. Together, these findings demonstrate that Intron P functions as a key regulatory element controlling *tim* mRNA nuclear export and, consequently, cytoplasmic mRNA and protein levels. These results suggest that the ratio of fully spliced to partially spliced *tim* transcripts is tightly regulated to generate a ∼24-hour circadian period and robust behavioral rhythms.

### Intron P is necessary and sufficient for nuclear retention of transcripts in Drosophila and human cells

We next asked whether Intron P alone is sufficient to drive nuclear retention of any transcript. To this end, we generated two UAS transgenic fly lines: an ‘IntP’ line in which Intron P and its flanking exons were inserted into a mCherry NLS TetR reporter (Fig. 3A), and an ‘Exon’ control line containing only the flanking exons (Fig. 3A). The flanking exonic sequences were included to control for potential effects of exonic splicing enhancers and silencers ^26^. To ensure similar rates of transcription for both these reporters, they were inserted into the same genomic location (see Methods). We drove reporter expression in clock neurons using the *Clk-GAL4* driver and visualized reporter transcripts using TetR-targeted FISH probes (see Methods). In ‘Exon’ control flies, reporter transcripts were predominantly cytoplasmic, with only ∼25% detected in the nucleus (Fig. 3B, 3C). This nuclear fraction likely represents transcripts in the process of being exported. In contrast, in ‘IntP’ flies, reporter transcripts were largely retained in the nucleus (∼75% nuclear; Fig. 3B, 3C), demonstrating that Intron P alone is sufficient to drive nuclear retention of transcripts.

**Fig. 3.**
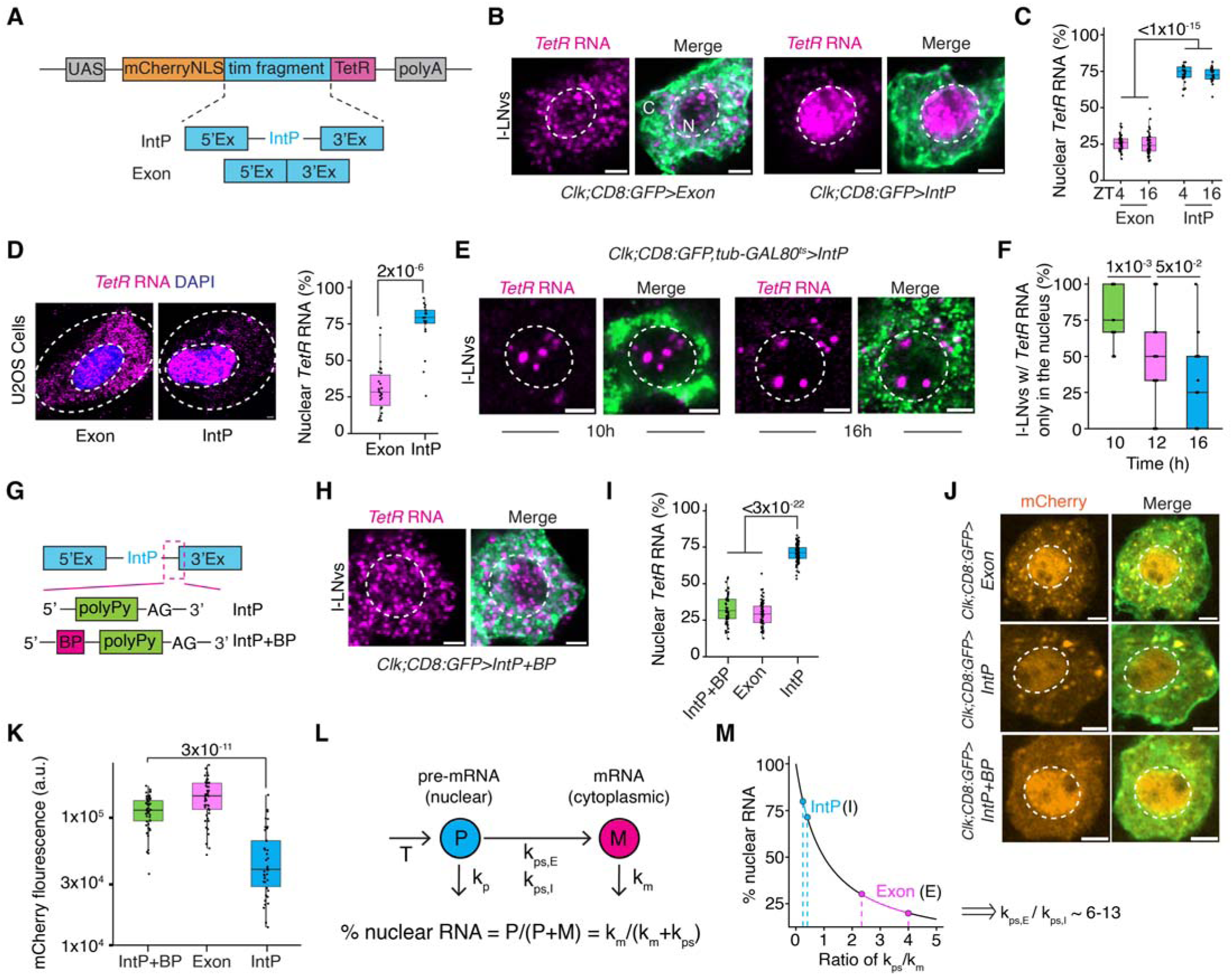
**Inefficient splicing of Intron P promotes nuclear retention of reporter transcripts in both *Drosophila* neurons and human cells**. (**A**) Schematic of UAS reporter fly lines containing Intron P and its flanking exons (IntP), or only the flanking exons without Intron P (Exon). (**B**) Representative images of *tetR* RNA in l-LNvs from Exon and IntP fly lines. l-LNv cytoplasm is shown in green, *tetR* transcripts are shown in magenta, and nuclei are outlined in white. N denotes nucleus, C denotes cytoplasm. (**C**) Quantification of the percentage of nuclear *tetR* RNA in l-LNvs from Exon (pink) and IntP (blue) flies at different time points (ZT4, ZT16) across the circadian cycle. (**D**) Representative images of *tetR* RNA (magenta) in U2OS cells transfected with Exon (left) and IntP (right) reporters. Nuclei are stained with DAPI (blue) and outlined in white. Quantification of the percentage of nuclear *tetR* RNA in U2OS cells is shown in the right panel. (**E**) Representative images of *tetR* RNA in l-LNvs from IntP flies at 10 and 16 hours after flies were shifted to 29 °C to induce *tetR* RNA expression. (**F**) Quantification of percentage of l-LNvs which possess *tetR* RNA exclusively in the nucleus at different time points (10 h, 12 h, 16 h) after the temperature shift to 29 °C. (**G**) Schematic of a UAS reporter fly line containing Intron P with a consensus branch point sequence inserted near its poly-pyrimidine tract (IntP+BP). (**H**) Representative image of *tetR* RNA (magenta) in a l-LNv from IntP+BP flies. (**I**) Quantification of the percentage of nuclear *tetR* RNA in l-LNvs from IntP+BP (green), Exon (pink), and IntP (blue) flies. (**J**) Representative images of mCherry protein in l-LNvs from Exon (top), IntP (middle), and IntP+BP (bottom) flies. l-LNv cytoplasm is shown in green, and mCherry protein is shown in orange. (**K**) Quantification of mCherry protein fluorescence intensity per l-LNv from IntP+BP (green), Exon (pink), and IntP (blue) flies. (**L**) Schematic of a two-state kinetics model for *tetR* RNA processing. Under steady-state conditions, a constant transcription rate produces nuclear pre-mRNAs containing Intron P (P), which are processed into mature mRNAs and exported to the cytoplasm (M). Both pre-mRNAs (P) and mature mRNAs (M) are subject to constant-rate degradation in the nucleus and cytoplasm, respectively (see Methods for further details). ‘T’ refers to the transcription rate, ‘k_p_’ refers to the nuclear mRNA degradation rate, ‘k_m_’ refers to the cytoplasmic mRNA degradation rate, ‘k_ps,E_’ refers to pre-processing rate of *TetR* reporter transcripts (which do not contain Intron P) from the UAS-Exon fly line, ‘k_ps,I_’ refers to the pre-processing rate of *TetR* reporter transcripts (which contain Intron P) from the UAS-IntP fly line. (**M**) Schematic plot illustrating how inefficient processing of pre-mRNA (P) can lead to nuclear enrichment of RNA. Experimental data indicate that the nuclear fraction of *tetR* RNA (*i.e.*, percentage of nuclear *tetR* RNA; see first and third quartiles in panel 3c) ranges from 21–29% in the Exon reporter flies (pink lines) and 70–77% in the IntP reporter flies (blue lines). These measurements suggest that RNA processing is ∼6 to 13-fold less efficient when Intron P is present. In **C**, **D**, **F**, **I, and K,** data are presented as box plots displaying the median and quantiles. Statistical analysis was performed using two-sided Mann-Whitney U tests. In all plots, each dot represents data from l-LNvs of a single *Drosophila* hemi-brain. Refer to Supplementary Table S10 for all statistical comparisons. l-LNv nuclei are circled in white. Scale bars-2µm.

To determine whether this mechanism is conserved across cell types, we expressed the IntP reporter in *Drosophila* courtship neurons using the *Fru-GAL4* driver (see Methods). Because these neurons lack endogenous tim expression, all TetR and Intron P signals originate solely from the reporter. In courtship neurons, ∼70% of TetR transcripts were nuclear—similar to the 75% observed in clock neurons—with nuclear TetR signals colocalizing with Intron P and no Intron P signal detected in the cytoplasm (Extended Data Fig. 8A, 8E). These results demonstrate that Intron P-containing transcripts are retained in the nucleus and only fully spliced transcripts are exported. We next asked whether Intron P’s nuclear retention function is conserved in human cells by transiently overexpressing IntP and Exon reporters in the human osteosarcoma cell line U2OS. Reporter transcripts in IntP-transfected cells were significantly more nuclear compared to those in Exon-transfected controls (Fig. 3D), suggesting that Intron P-mediated retention is conserved across species. Notably, total reporter transcript levels were slightly higher in IntP versus Exon lines (Extended Data Fig. 8C), possibly due to increased nuclear stability or enhanced transcription driven by Intron P.

We next investigated the splicing dynamics of Intron P-containing reporter transcripts (*TetR* mRNAs) from the ‘IntP’ line by tracking their subcellular localization over time. Since transcripts retaining Intron P remain nuclear while fully spliced transcripts are exported, subcellular localization provides a clear, single-cell-level indicator of splicing status. To induce *TetR* mRNA expression, we used the *Clk GAL4, tub GAL80^ts^* system, which enables temperature controlled activation of the UAS transgenes (see Methods). We shifted adult flies to the permissive temperature (29 °C) to trigger *TetR* mRNA production, and performed RNA-FISH experiments at three timepoints, 10h, 12h, and 16h post-temperature shift. At 10 hours, the earliest time point when *TetR* transcripts were detectable, the transcripts were almost entirely nuclear (Fig. 3E, 3F). This is consistent with our earlier observation that ∼90% of newly synthesized *tim* mRNAs retain Intron P (Fig. 1H). By 12 and 16 hours post-temperature shift, however, *TetR* mRNAs were detected in both the nucleus and cytoplasm (Fig. 3E, 3F; Extended Data Fig. 8b), suggesting that Intron P had been spliced out in a subset of transcripts, enabling their export. Together, these results demonstrate that Intron P is both necessary and sufficient for nuclear retention and functions as a molecular “tag” for mRNA retention across species.

### Inefficient splicing of Intron P drives nuclear mRNA retention

Given that Intron P has strong donor and acceptor splice sites but lacks a canonical branch point, a sequence essential for efficient splicing ^27^, we investigated whether inserting a branch point sequence would improve its splicing efficiency. To this end, we created a new UAS transgenic fly line, ‘IntP+BP’, by adding a consensus branch point (BP) sequence to the

IntP reporter (Fig. 3G). Unlike the ‘IntP’ line, where reporter transcripts were predominantly nuclear, transcripts from ‘IntP+BP’ line were mostly cytoplasmic, resembling the Exon control (which lacks Intron P), in both *Drosophila* clock neurons (Fig. 3H, 3I; Extended Data Fig. 8D-E) and human U2OS cells (Extended Data Fig. 8F), indicating that adding a branch point sequence boosts splicing efficiency. These results unambiguously show that inefficient splicing of Intron P, rather than other mechanisms such as nuclear export, governs mRNA nuclear retention. Moreover, enhancing splicing efficiency—whether through branch-point insertion or intron removal—promotes nuclear export. To assess functional consequences of mRNA nuclear retention, we measured mCherry fluorescence in clock neurons from all three reporter lines. The ‘IntP’ line exhibited significantly lower signal compared to the ‘Exon’ and ‘IntP+BP’ lines (Fig. 3J, 3K), consistent with reduced cytoplasmic mRNA available for translation in ‘IntP’ lines.

To quantify Intron P’s splicing efficiency, we adapted a kinetic model ^28^ that estimates splicing rates based on the steady-state nuclear versus cytoplasmic distribution of reporter transcripts (see Methods, Fig. 3L). Under steady-state conditions, the rates of transcription, export, and mRNA degradation are constant and similar for the ‘IntP’ and ‘Exon’ reporters. The ‘Exon’ reporter, which lacks introns, serves as a control representing fully processed transcripts. In the ‘IntP’ condition, transcripts containing Intron P are retained in the nucleus until fully spliced (Fig. 3L); therefore, the ratio of nuclear to cytoplasmic transcripts reflects the splicing efficiency of Intron P—higher nuclear retention indicates less efficient splicing. Applying the model to data from the ‘IntP’ and ‘Exon’ reporter assays, we estimated that Intron P-containing transcripts are processed ∼10-fold less efficiently than exon-only transcripts (see Methods, Fig. 3M).

Next, we investigated whether environmental factors affect the splicing efficiency of Intron P by using the ‘IntP’ reporter line. While splicing efficiency did not change significantly across the light-dark cycle (Extended Data Fig. 8G-I), it increased modestly at lower temperatures, as evidenced by a slightly larger fraction of *TetR* transcripts localizing to the cytoplasm (Extended Data Fig. 8J-L). Together, these results demonstrate that inefficient splicing of intron P drives nuclear retention, while enhancing splicing efficiency promotes cytoplasmic export and increases protein production.

### Knockdown of spliceosome factors affects Intron P splicing

Next, we investigated the molecular mechanisms that regulate the splicing of *tim* Intron P. Splicing is carried out by the spliceosome ^29^, a large protein-RNA complex composed of small nuclear ribonucleoproteins (snRNPs), including U1, U2, the U4/U6.U5 tri-snRNP, and the Prp19 complex. To identify factors specifically regulating intron P splicing, we first performed an RNAi screen targeting 45 spliceosome components by knocking them down in clock neurons and assaying their behavioral rhythms. We next focused on a subset of candidates that produced strong circadian rhythm defects and analyzed the effects of their knockdown using RNA-FISH and transcriptomic analyses to determine whether they affect splicing of Intron P (Fig. 4A; Supplementary Tables S3, S8). First, we examined Precursor RNA Processing 3 (Prp3), a U4/U6.U5 tri-snRNP component ^30, 31^, as clock neuron specific knockdown of Prp3 (∼50% knockdown efficiency; Extended Data Fig. 9F) completely abolished rhythmic behavior (Fig. 4B; Supplementary Tables S3, S8). RNA-FISH experiments revealed that knockdown of Prp3 in clock neurons significantly increased the nuclear retention of *tim* mRNAs containing Intron P, from ∼50% in controls to ∼80% in *Prp3-RNAi* flies (Fig. 4C-E; Extended Data Fig. 9A). Strikingly, the splicing of other *tim* introns was unaffected (Fig. 4F, 4G), indicating that Intron P splicing is selectively impaired upon ∼50% knockdown of Prp3. As a result, cytoplasmic levels of fully spliced *tim* mRNA and TIM protein levels were significantly reduced throughout the circadian cycle (Fig. 4H, 4I), which disrupts clock function and leads to arrhythmic behavior. Expression of an siRNA □resistant Prp3 transgene in *Prp3*□*RNAi* flies rescued the wild-type-like∼50% nuclear-50% cytoplasmic distribution of *tim* mRNAs and restored behavioral rhythmicity (Extended Data Fig. □9B-E). Together, these experiments demonstrate Prp3’s essential role in intron P splicing.

**Fig. 4.**
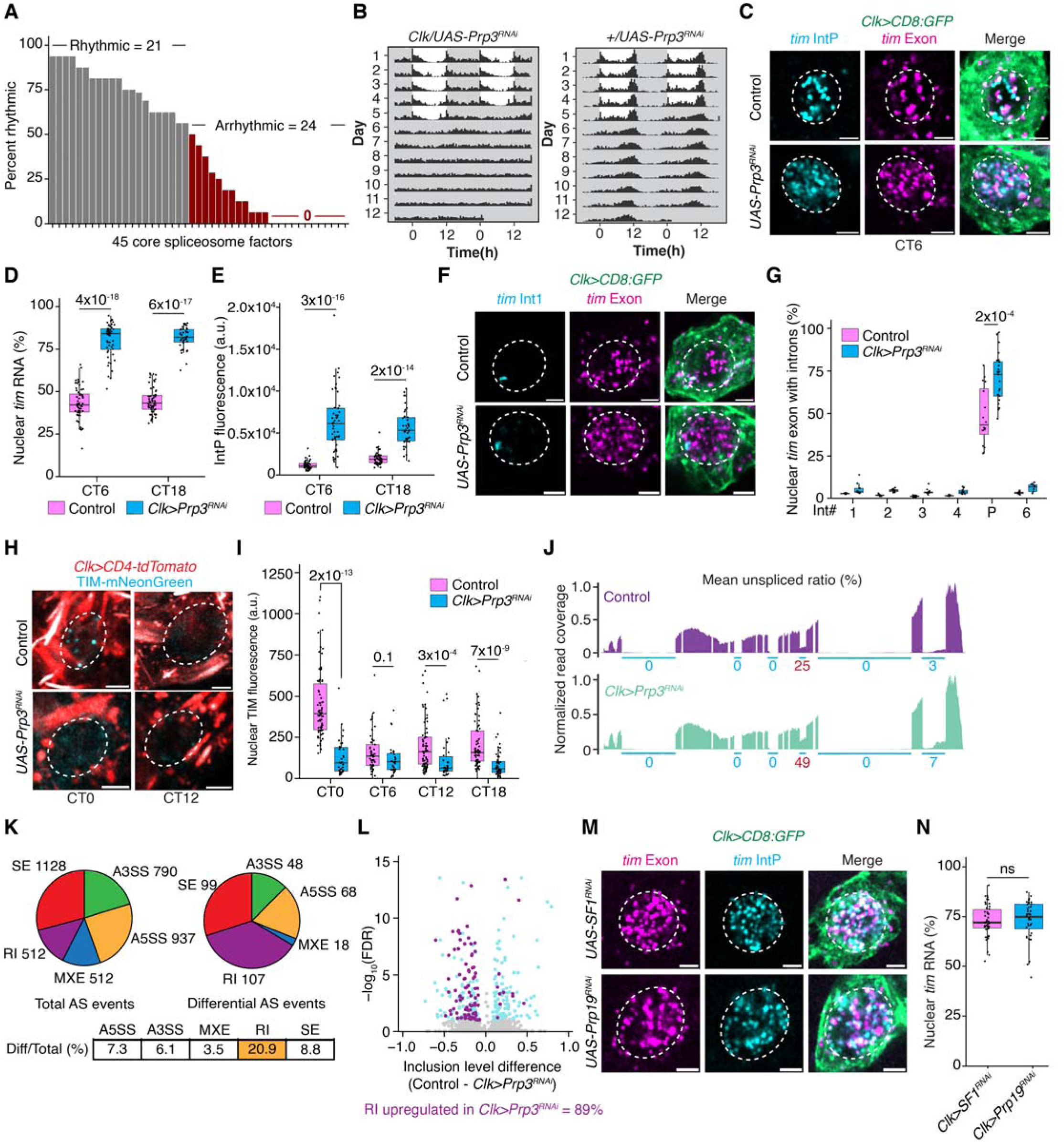
Intron P splicing is affected by knockdown of spliceosome factors. (**A**) RNAi-mediated knockdown of 45 spliceosome factors was performed in clock neurons to assess their impact on circadian locomotor rhythms. Rhythmicity was evaluated under constant conditions (DD), and knockdowns were classified as “arrhythmic” if fewer than 50% of the flies exhibited rhythmic behavior (Supplementary Tables S3, S8). (**B**) Averaged population locomotor-activity profiles of *Clk>Prp3-RNAi* flies (left panel) and control flies (right panel) in LD and DD conditions, with rest-activity patterns shown for two consecutive days in each row. *Clk>Prp3-RNAi* flies display arrhythmic behavior in DD (Supplementary Tables 3, 8). (**C**) Representative images of *tim* Intron P (cyan) and exon (magenta) FISH signals in l-LNv clock neurons from *Clk>Prp3-RNAi* and control flies. (**D, E**) Quantification of percentage of nuclear *tim* RNA (**D**) and Intron P fluorescence intensity (**E**) in l-LNvs from *Clk>Prp3-RNAi* and control flies. Compared to the wildtype control, knockdown of Prp3 leads to increased nuclear retention of *tim* RNA and higher levels of Intron P in the nucleus. (**F**) Representative images of *tim* Intron 1 (cyan) and exon (magenta) FISH signals in l-LNvs from *Clk>Prp3-RNAi* and control flies. (**G**) Percentage of nuclear *tim* exon spots colocalizing with the intron spots in l-LNvs from *Clk>Prp3-RNAi* and control flies. (**H**) Representative images of endogenous TIM-mNG protein in l-LNvs from *Clk>Prp3-RNAi* and control flies during the repression (CT0) and the activation (CT12) phases. (**I**) Quantification of TIM-mNG protein fluorescence intensity in l-LNvs over the circadian cycle. (**J**) Normalized read coverage of the *tim* locus from poly(A)-seq in Prp3-RNAi knockdown (top) and control samples (bottom). Splicing status of *tim* introns, quantified by mean unspliced read ratio (averaged over three replicates) is shown as percentages. Intron P unspliced ratio is marked in red (Supplementary Table S4). (**K**) Pie chart summarizing the differential analysis of alternative splicing (AS) events between Clk>Prp3-RNAi and control samples (see Supplementary Table S5). (**L**) Volcano plot of differential AS events, with significant events shown in cyan and significant retained intron events in purple (see Methods). Among the differentially retained introns, Prp3 knockdown caused increased intron retention a majority of the times (89%). (**M**) Representative images of *tim* exon (magenta) and Intron P (cyan) FISH signals in l-LNvs at CT0 under Prp19-RNAi and SF1-RNAi knockdown conditions. (**N**) Quantification of the percentage of nuclear *tim* transcripts in l-LNvs under Prp19 and SF1 knockdown conditions. In **D**, **E**, **G**, **I, and N,** data are presented as box plots displaying the median and quantiles. Statistical analysis was performed using two-sided Mann-Whitney U tests. In all plots, each dot represents data from l-LNvs of a single *Drosophila* hemi-brain. Refer to Supplementary Table S10 for all statistical comparisons. l-LNv nuclei are circled in white. Scale bars-2µm.

To evaluate the broader role of Prp3 in splicing, we performed RNA-sequencing on FACS-sorted clock neurons from Prp3-RNAi and control flies (Extended Data Fig. 9F). The analysis revealed that Prp3 knockdown specifically increased retention of the *tim* intron P, while the splicing of all other *tim* introns, which were co-transcriptionally spliced, remained unaffected (Fig. 4J; Supplementary Table S4), consistent with our RNA-FISH results. Extending this observation genome-wide, we found that an overwhelming majority (99.4%) of constitutively spliced introns (∼26,000 total) showed no changes upon Prp3 depletion (Fig. 9G; Supplementary Table S4). Notably, among introns that are naturally retained in wild-type cells, only about 5% exhibited further increased retention after Prp3 knockdown, mirroring the effect seen with *tim* intron P (Fig. 9G; Supplementary Table S4). We then examined how Prp3 depletion influenced alternative splicing events, and found that retained introns were the most significantly affected (Fig. 4K-L; Supplementary Table S5). Importantly, these Prp3-sensitive introns typically possessed weaker 5’ and 3’ splice sites or branchpoints and are more evolutionarily conserved (Extended Data Fig. 9H, 9I), suggesting that Prp3 plays a crucial role in facilitating the splicing of introns with suboptimal splicing signals. These results are consistent with prior findings in yeast, where the Prp4 kinase, another tri-snRNP component, is specifically required for the efficient splicing of introns with weak splice sites and branchpoints ^32^.

To explore whether intron P splicing depends on additional spliceosome factors, we knocked down Splicing Factor 1 (SF1), which binds the branch point sequence ^27^, and Prp19, a core component of the Prp19 complex ^33^, in clock neurons. Both knockdowns resulted in similar intron P retention defects and behavioral arrhythmicity (Fig. 4M, 4N; Extended Data Fig. 9J; Supplementary Tables S3, S8).

### A genetic screen to identify *tim* Intron P splicing activators and repressors

We next investigated whether specific RNA binding proteins (RBPs) regulate the splicing of *tim* Intron P. RBPs influence splicing by interacting with specific RNA sequences or structures, modulating the assembly or activity of the spliceosome to either enhance or repress splicing ^34^. To identify potential regulators of *tim* Intron P splicing, we conducted a behavioral RNAi screen by systematically knocking down 37 RBPs in clock neurons (Fig. 5A; Supplementary Tables S7, S8). These RBPs were previously shown to affect splicing in *Drosophila* S2 cells ^34^. For RBPs whose knockdown disrupted rhythmic behavior, we used RNA-FISH assays to examine the effects of their knockdown on *tim* Intron P splicing and mRNA localization. Strikingly, clock neuron-specific knockdown of Qkr58E□2, a *Drosophila* homolog of the human KH□domain RNA binding protein SAM68 ^35^, caused strong arrhythmicity (Fig. 5B). SAM68 has been shown to regulate the splicing of introns of many key genes such as SMN2 ^36^, BCLx ^37^, Cyclin D1 ^38^, and mTor ^39^, acting as either an activator or repressor depending on the context. For instance, in the *mTor* gene, SAM68 was shown to bind to the U(U/A)AA motif within a specific intron and recruit U1 snRNP to the 5’ splice site, thereby enhancing splicing efficiency ^39^.

**Fig. 5.**
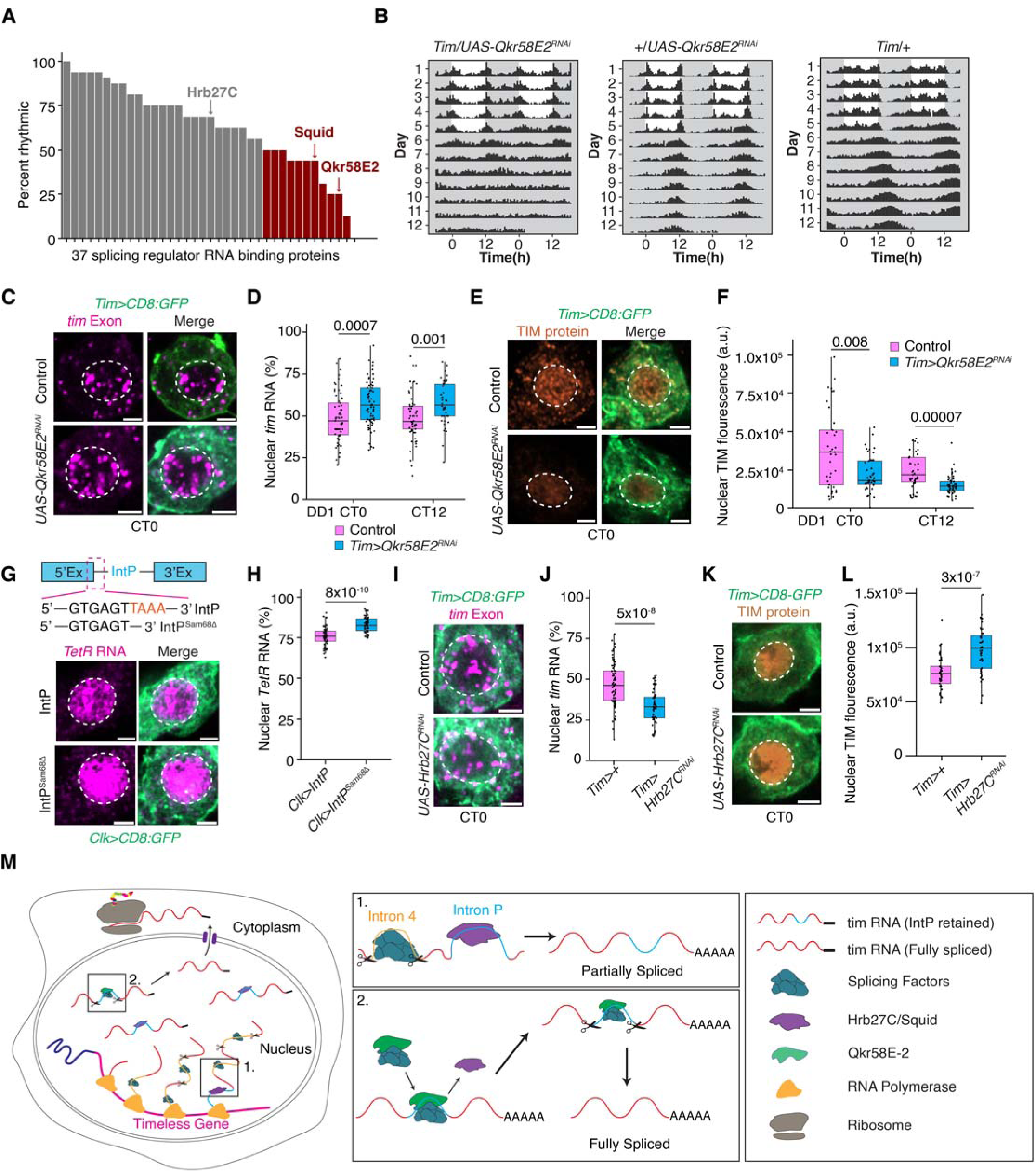
Intron P splicing is activated by Qkr58E-2 and repressed by Hrb27C and Squid. (**A**) RNAi-mediated knockdown of 45 spliceosome factors was performed in clock neurons to assess their impact on circadian locomotor rhythms. Rhythmicity was evaluated under constant conditions (DD), and knockdowns were classified as “arrhythmic” if fewer than 50% of the flies exhibited rhythmic behavior (Supplementary Tables S7, S8). (**B**) Averaged population locomotor-activity profiles of *Clk>Qkr58E2-RNAi* flies (left panel) and control flies (right panels) in LD and DD conditions, with rest-activity patterns shown for two consecutive days in each row. *Clk>Qkr58E2-RNAi* flies display arrhythmic behavior in DD (Supplementary Tables 7, 8). (**C**) Representative images of *tim* exon spots (magenta) in l-LNvs under *Clk>Qkr58E2-RNAi* and control conditions. (**D**) Quantification of the percentage of nuclear *tim* RNA in l-LNvs under *Clk>Qkr58E2-RNAi* and control conditions. Knockdown of Qkr58E-2 leads to increased nuclear retention of *tim* RNA. (**E**) Representative images of TIM protein (orange) in l-LNvs under *Clk>Qkr58E2-RNAi* and control conditions. (**F**) Quantification of TIM protein fluorescence intensity in l-LNvs at repression (CT0) and activation (CT12) phases. (**G**) Schematic showing predicted Qkr58E2/Sam68 binding site (UAAA) near the 5’ splice site of Intron P and the design of a TetR reporter line where this motif is deleted (IntP^Sam68Δ^, top). Representative images of *TetR* RNA (magenta) in l-LNvs from the control (IntP) and motif-deleted (IntP^Sam68Δ^) reporter fly lines. (**H**) Quantification of percentage of nuclear *TetR* RNA in l-LNvs from IntP and IntP^Sam68Δ^ flies. (**I-L**) Representative images of *tim* RNA (**I**), quantification of percentage of nuclear *tim* RNA (**J**), representative images of TIM protein (**K**), and quantification of nuclear TIM protein fluorescence (**l**) in l-LNvs under *Tim>Hrb27C-RNAi* and control conditions. (**M**) Model of post-transcriptional splicing of *tim* RNAs. Intron P is retained in nearly all nascent *tim* transcripts, is spliced inefficiently post transcriptionally, and only fully spliced transcripts are exported. In **D**, **F**, **H**, **J, and L,** data are presented as box plots displaying the median and quantiles. Statistical analysis was performed using two-sided Mann-Whitney U tests. In all plots, each dot represents data from l-LNvs of a single *Drosophila* hemi-brain. Refer to Supplementary Table S10 for all statistical comparisons. l-LNv nuclei are circled in white. Scale bars-2µm.

In our studies, knockdown of Qkr58E 2 in clock neurons significantly increased nuclear retention of *tim* mRNA (Fig. 5C, 5D) and decreased TIM protein levels (Fig. 5E, 5F), suggesting that Qkr58E 2 functions as an activator of Intron P splicing. To further evaluate the effect of Qkr58E-2 on Intron P splicing, we utilized our intron reporter system to monitor splicing efficiency *via* transcript localization, as described in Fig. 3a. We identified a U/A-rich binding motif near Intron P’s 5’ splice site, similar to the motif recognized by SAM68 ^40^. To test the functional role of this motif, we generated a UAS transgenic fly line, *IntP^Sam68^*^Δ^ by deleting the U/A-rich motif from the Intron P sequence (Fig. 5G). Compared to the control IntP line, the *IntP^Sam68^*^Δ^ line showed significantly increased nuclear retention of *TetR* reporter transcripts (Fig. 5G, 5H; Extended Data Fig. 10B), suggesting that Intron P splicing is impaired in the ‘IntP-Sam68Δ’ line. This supports a model in which Qkr58E 2 binds to the U/A rich motif near the 5′ splice site of Intron P, thereby promoting its splicing. To explore Qkr58E-2’s broader impact on splicing, we analyzed a previously published RNA-seq dataset from *Drosophila* S2 cells ^34^ and found that knockdown of Qkr58E-2 affected various alternative splicing events, including intron retention, indicating that Qkr58E-2 serves as a general regulator of splicing (Extended Data Fig. 10A).

In addition to Qkr58E-2, our screen also identified two hnRNP proteins, Hrb27C and Squid, which can act as splicing repressors ^41, 42^—whose knockdown in clock neurons also affected circadian rhythmic behavior (Supplementary Table S7). Knockdown of either factor significantly decreased nuclear retention of *tim* mRNA, indicating more efficient removal of intron P and export of fully spliced transcripts (Fig. 5I, 5J; Extended Data Fig. 10C, 10D). Consistent with increased cytoplasmic *tim* mRNA, TIM protein levels were elevated upon knockdown of Hrb27C or Squid compared to controls (Fig. 5K, 5L; Extended Data Fig. 10E-F). Together, these data suggest that Hrb27C and Squid repress intron P splicing. Interestingly, knockdown of several other RBPs also disrupted rhythmicity, but FISH experiments revealed no effect on the nuclear–cytoplasmic distribution of *tim* mRNAs (Extended Data Fig. 10G-I; Supplementary Table S7). This suggests that while Qkr58E 2, Hrb27C, and Squid modulate circadian rhythms through *tim* splicing, these additional RBPs might influence the splicing of other transcripts essential for circadian rhythms.

Together, these results reveal a whole new level of post-transcriptional regulatory mechanism in clock neurons: Qkr58E-2 activates Intron P splicing, promoting *tim* mRNA export, while Hrb27C and Squid repress Intron P splicing, driving nuclear retention. Our model proposes that repressors are initially recruited to nascent *tim* mRNA to suppress intron P splicing, followed by the recruitment of activators, such as Qkr58E-2, to promote splicing (Fig. 5M). This interplay precisely regulates intron P splicing efficiency, thereby controlling the amount of translatable tim mRNA in the cytoplasm and the resulting TIM protein levels to maintain robust circadian rhythms.

## Discussion

Our findings uncover a previously unrecognized RNA-based delay mechanism—a molecular “timer”—in which inefficient post-transcriptional splicing of a single intron, Intron P, within the *timeless* (*tim*) gene plays a central role in establishing the ∼24-hour circadian period. While previous studies have identified protein-based delay mechanisms in the circadian clock, such as phosphorylation and proteasomal degradation of the repressors PER ^3, 4^ and TIM ^5^, or the slow ATPase activity of KaiC ^6^ and RUVBL ^7^ proteins, our work, in contrast, reveals a novel post-transcriptional rate-limiting step that acts upstream of nuclear export. Using single-molecule imaging, nascent RNA-seq, and computational modeling, we unexpectedly found that that Intron P is consistently bypassed by the spliceosome during transcription due to its weak splicing signals and is instead spliced post-transcriptionally with ∼10-fold lower efficiency. This gives rise to two populations of *tim* transcripts: unspliced RNAs containing Intron P that are retained in the nucleus, and fully spliced transcripts that are exported to the cytoplasm. Importantly, unlike typical alternative splicing events that generate protein isoform diversity ^8^, Intron P retention creates a nuclear export bottleneck that fine-tunes TIM protein levels without altering the protein’s isoform.

Inefficient splicing of Intron P, far from being a defective RNA processing event, serves as a crucial regulating mechanism regulating TIM protein levels and circadian timing in *Drosophila*. CRISPR-mediated removal of Intron P abolishes nuclear retention, accelerates TIM accumulation, advances the circadian phase by ∼2 hours, and reduces behavioral rhythmicity to ∼60%. Furthermore, Intron P is evolutionarily conserved across all *Drosophila* species, with its inefficient splicing and nuclear localization similarly conserved. Remarkably, Intron P alone is sufficient to induce nuclear retention of reporter mRNAs in both *Drosophila* and human cells, suggesting its broader potential as a tool for spatial regulation of mRNA. To better understand the basis of this regulatory function, we identified multiple layers of control governing intron P splicing. Its inefficiency arises from intrinsic sequence features—most notably, the absence of a canonical branch point sequence—and is further regulated by trans-acting factors. These include the splicing activator Qkr58E-2 (a Sam68 homolog) and the repressors Hrb27C and Squid (hnRNP proteins), which collectively modulate the balance between spliced and unspliced *tim* transcripts—thereby positioning intron P as a molecular timer that governs circadian period and amplitude.

Intriguingly, a recent transcriptomic study revealed that *Per1* mRNA is evenly distributed between the nucleus and cytoplasm in mouse liver cells ^43^. Based on intron P’s role in our study, we propose that inefficient splicing–mediated nuclear retention may also be a conserved mechanism in mammalian circadian systems. Beyond circadian rhythms, we identified 75 additional introns in *Drosophila* brain transcripts that exhibit splicing dynamics similar to Intron P, including in genes such as *unc13*, a synaptic vesicle release regulator and ALS risk factor gene. More broadly, nuclear retention of unspliced, polyadenylated transcripts is widespread across eukaryotes, including humans, mice, and plants ^10–12^, indicating that mRNA processing delays may constitute a general principle of gene control in diverse biological contexts, such as synaptic function, neural activity, and development.

Together, our results reveal how modulating the timing and efficiency of a single intron of a core clock gene introduces regulatory delays in circadian rhythms, providing a flexible molecular strategy for shaping protein dynamics across biological systems. The *timeless* Intron P exemplifies how evolution can repurpose splicing inefficiency into a functional feature, transforming a passive intronic sequence into an active temporal gate for gene expression.

## Methods

### Fly stocks

Flies were raised on standard cornmeal/yeast media and maintained at 25C with 50-70% humidity under a 12h:12h Light-Dark (LD) schedule. Flies were entrained to Light-Dark (LD) cycles where they were exposed to 12-hour Light-Dark (LD) cycles for 5-7 days, followed by a shift to complete darkness (DD) for another 7 days. The initiation of the light phase is labeled as Zeitgeber Time (ZT) 0, when lights are turned on, while ZT12 is the beginning of the dark period, when lights are off. When referencing the times in the continuous darkness phase, we use Circadian Time (CT). CT0 (subjective dawn) is the time when lights would have been turned on and CT12 (subjective dusk) is the time when lights would have been off. Labels DD1 and DD2 represent the first and second days of complete darkness. The following flies used in the study were previously described or obtained from the Bloomington Stock Center: *w1118* (BL-6326), *y,sc,v* (BL-25709), *CLK-GAL4* (BL-93198), *Fru-GAL4* (BL-66696), *tub-gal80^ts^* (BL-7019), *UAS-Tim-GAL4* (BL-80941), *UAS-Dcr2* (BL-24644), *per^01^* (BL-80917), *UAS-mCD8GFP* (BL-32184), *UAS-CD4tdTom* (BL-35841). Specific genotypes of flies used in the experiments are presented in the figures and in the Supplementary Tables S3, S7, and S8.

### Design of new transgenic fly lines

The following fly lines are created for this study: ;tim-IntPΔ; (CRISPR knockout of Intron P), UAS-Exon, UAS-IntP, UAS-IntP+BP, UAS-IntP_Sam68Δ. Refer to Supplementary Table S9 for exact plasmid sequences. To make the CRISPR knockout fly line, we followed a previously established protocol ^44, 45^. In brief, two gRNAs GTTCACTGGCGTCGAACTTG (gRNA-1) and GAACCTTGAGACCACTCCGC (gRNA-2) where cloned into the gRNA expression plasmid pU6-BbsI-chiRNA (Addgene #45946) using restriction enzyme (BbsI) cloning, which were confirmed by Sanger sequencing with T7 primer and prepared using Midi-prep (QIAGEN). Donor plasmid was cloned with the pBlueScriptII SK(-) backbone (GenScript, Inc.) which contains complete flanking exons of intron P with PAM sequences mutated to maintain the coding amino acid sequence (gRNA-1 PAM mutated to AGA and gRNA-2 PAM mutated to AGC). gRNA and donor plasmids were microinjected into germline Cas9 expression embryos (nos-Cas9 attp2, Rainbow Transgenic Flies, Inc.) and crossed individually with chr2 balancer line. Successful knockout was screened by PCR, backcrossed to chr2 balancer line and homozygosed. Finally, correct editing was confirmed by Sanger sequencing and HCR RNA-FISH showing lack of intron P staining.

To make the UAS overexpression fly lines, Intron P, together with its immediate 5’ and 3’ flanking exons, are inserted in frame into a coding frame of mCherry-NLS-tetR between the NLS and tetR elements within a modified UAS expression plasmid pJFRC-MUH (Addgene #26213) where 3 UAS elements are kept for reduced overexpression strength. Flanking exons of Intron P encodes total of 209 amino acids. UAS-Exon only contains flanking exons of Intron P, UAS-IntP+BP has insertion of a canonical branch point sequence (TACTAAT) 5 base pairs upstream of the polypyrimidine tract CCTCTCTCCTTCTCCTCCTTCT. Refer to Supplementary Table S9 for exact plasmid sequences. Plasmids were cloned from gBlock fragments (IDT, Inc.) using Gibson assembly (NEBuilder HiFi DNA Assembly Master Mix) following manufacturer protocol. Clones were verified by whole plasmid sequencing (Eurofins, Inc.) and prepared for injection (QIAGEN midi-prep). Cloned plasmids were microinjected into phiC31 integrase line with docking site attP40 (Rainbow Transgenic Flies, Inc.)

We used the TRiP RNAi line (BDSC #55256) for targeted knockdown of Prp3. Exact RNAi targeting sequence was fetched from UP-TORR (https://www.flyrnai.org/up-torr/). The Prp3-RNAi line (Reagent ID HMC03943) target is ACGACGCAAGCGGTAAGATAA, which resides in the coding sequence (CDS) of Prp3. The targeting sequence was mutated extensively and synonymously to ATGATGCCTCGGGCAAAATTT (i.e., Prp3 protein sequence unchanged). The mutated CDS, together with a 3xV5 tag fused to the C terminal with a GGGS linker, was cloned into pJFRC-MUH (Addgene #26213) to generate the UAS overexpression construct by Gibson assembly. Clones were verified by whole plasmid sequencing (Eurofins, Inc.) and prepared for injection (QIAGEN midi-prep). Refer to Supplementary Table S9 for exact plasmid sequences and cloning strategy. Verified plasmid was microinjected into phiC31 integrase line with docking site attP2 (Line R8622, Rainbow Transgenic Flies, Inc.).

*CLK-GAL4, tub-GAL80^ts^* system^46, 47^ was used for temperature-dependent expression of the UAS-IntP reporter. We crossed IntP flies with *CLK-GAL4, tub-GAL80^ts^* flies, maintaining progeny at 18°C to suppress TetR mRNA transcription. After shifting adult flies to 29°C to induce reporter transcription, we conducted a time series of FISH experiments earliest 8-hours post-temperature shift using the tetR probe. 10-hour mark is the earliest time point where TetR mRNAs were detectable.

### Human U2OS cell culture reporter assay

We overexpressed the mCherry-NLS-tetR reporters (Exon, IntP, IntP+BP) in human U2OS cells following standard transient transfection protocol using pFUGW expression plasmid. Refer to Supplementary Table S9 for exact plasmid sequences. pFUGW_MCS1 backbone was a gift from the Sami Barmada lab. We used digestion cloning (enzymes HpaI + AgeI) to insert the reporter constructs (as described above for the UAS overexpression), which was verified by whole plasmid sequencing (Eurofins, Inc.) and prepared by QIAGEN Midi-prep. Refer to Supplementary Table S9 for exact plasmid sequences and cloning strategy. Cells were cultured in DMEM (Gibco 11965092) supplemented with 10% FBS (Corning 35016CV), 1% Pen-Strep (Gibco 15070063) and 1% Glutamax (Gibco 35050061). We maintained cells in 10cm culture plates (Falcon 353003) and passaged once cells reach ∼80% confluency using 1:3∼1:10 split ratios. To perform overexpression assay, we prepared coverslips coated with Poly-D-Lysine. Coverslips (No 1.5, Harvard Apparatus 1217N78) were rinsed in 70% ethanol, air dried, and sterilized with UV light in the culture hood. Subsequently, coverslips were submerged in 50 ug/mL Poly-D-Lysine solution (Sigma A-003-M) and nutated for 1-hour at room temperature for coating, rinsed three times with ultrapure water, and air dried. Subsequently, cells were split into 24-well plates (Corning 3524) with coated coverslips at the bottom of wells until ∼60% confluent.

Cells were transfected with Lipofectamine 2000 (Invitrogen 11668027) following manufacturer protocol and expression of reporter construct confirmed by mCherry fluorescence (typical transfection time ∼1-day). Subsequently, cells were rinsed gently two times in sterile PBS, fixed with 4% PFA for 10min at room temperature, and subjected to the standard HCR-FISH protocol as described below.

### RNA Fluorescence In Situ Hybridization (RNA-FISH)

We followed a previously established protocol for smFISH ^15^. We followed the following protocol for hairpin chain reaction FISH (HCR-FISH). In brief, brains were dissected in PBS buffer, fixed for 20min at room temperature in 4% PFA (diluted in PBS), washed with PBS buffer, and put in 70% ethanol overnight at 4C for permeabilization. Next day, brains were washed in FISH wash buffer (2xSSC, 10% formamide, 2mM RVC, 0.2% Tween-20) for 5min at room temperature, and subsequently hybridized at 37C overnight for a minimum of 16-hours (2xSSC, 10% formamide, 10% dextran sulfate, 2mM RVC, 0.2% Tween-20, 5nM probe). Hybridized brains were washed in FISH wash buffer two times for a total of 1-hour wash at 37C to remove unbound probes and subsequently mounted in Prolong Glass mountant (smFISH) or proceed with amplification (HCR-FISH) overnight for a minimum of 16-hours (2xSSC, 10% dextran sulfate, 2mM RVC, 0.2% Tween-20, 60nM of each amplifier hairpin). Amplified brains were washed in 2xSSC + 0.2% Tween-20 two times for a total of 1-hour before mounting. Probes and amplifiers were purchased from LGC Bioresearch (smFISH) or Molecular Instruments (HCR-FISH). Reagents purchased from other vendors are 20xSSC (Invitrogen AM9770), formamide (Sigma F9037), 200mM RVC (NEB S1402S), Tween-20 (Sigma P2287), dextran sulfate (Fisher 433241000), absolute ethanol (Fisher A409), Prolong Glass mountant (Invitrogen P36980), 10xPBS (Invitrogen AM9624), Ultrapure water (Invitrogen 10977), and 16% PFA (EMS 15710).

### Immunostaining

Brains were dissected in PBS buffer, fixed for 20min at room temperature in 4% PFA, and then rinsed three times and permeabilized for 1-hour (20min, 3 times) in PBS + 0.2% Triton X-100. Brains are then block in PBS + 0.2% Tween-20 + 5% NGS (blocking buffer) for 3-hour at room temperature and subsequently incubated in blocking buffer + primary antibody (1:400 Rb anti-TIM, 1:200 Ms anti-SC35, 1:200 Ms anti-Pdf). The incubation was over two nights at 4C. Brains were subsequently rinsed three times and washed for 1-hour (20min, 3 times) in PBS + 0.2% Tween-20 (PBSTw) and then exchanged into blocking buffer + secondary antibody (1:1000 for all experiments). Incubation was overnight at 4C. Brains were subsequently rinsed three times and washed for 1-hour (20min, 3 times) in PBSTw before mounting with Prolong Glass mountant. Rb anti-TIM was a gift from Joanna Chiu Lab and Ms anti-SC35 was sc-53518 (Santa Cruz Biotechnology). Ms anti-Pdf was C7 (Developmental Studies Hybridoma Bank). Reagents purchased from other vendors are Triton X-100 (Sigma T8787), NGS (MP Bio 191356), goat anti-Rabbit 647 (Fisher A32733), and goat anti-Mouse 488 (Fisher A11001).

### HCR-FISH data analysis

We followed a previously established protocol for analysis of the FISH data involving segmentation of clock neurons and detection of diffraction-limited spots^15^. To estimate the percentage of FISH spots in the nucleus, we developed a workflow where 2D regions of interest (ROIs) are drawn manually at center slice of clock neurons. Spots within the ROIs are defined by the following criteria: 1) they are located within the ROI or 5px (∼300nm) distance from the ROI, and 2) intensity is within top 50 percentile of all spots and extracted from the list of all detected diffraction-limited spots. To perform colocalization analysis between two FISH channels in the same image, we developed a custom function that computes all pairwise distances among spots in two channels. For each spot, the ‘nearest neighbor distance’ (NND) is defined as the smallest 3-D distance from this spot to all spots in the other channel. If the NND value is below a cutoff (400nm), the spot is classified as ‘colocalized to the other channel’. This then allows quantification of the percentage of spots that colocalizes with the other channel. Custom ImageJ plugin for ROI-based spot classification is available on Github (https://github.com/yeyuan98/punctaTracker) and ImageJ site for easy installation (https://sites.imagej.net/Yuanye1998/). Reproducible workflow for further processing of classified spots and nearest neighbor distance computation is available as a manual in R package ‘ijAnalysis’ (https://yeyuan98.r-universe.dev/articles/ijAnalysis/spotInRoi.html).

### RNA-seq library preparation and sequencing

Flies of genotype ClkGAL4>UAS-GFP-NLS was used for a bright GFP marker specific to clock neurons and DAPI was used as a viability marker. A small population of cells were heated at 60C for 5min and used as a dead cell (DAPI+) control. Live clock neurons are defined by GFP+ and DAPI-gating. For each sample, ∼2,000 live clock were sorted with FACS sorter (Sony MA900) directly into cell lysis buffer (NEBNext Low-Input cDNA Synthesis Module) in 1.5mL DNA low-bind tube (Eppendorf). Final volume of the lysis buffer (with sorted cells) is ∼16uL. Cells were lysed and half of the lysate was subsequently used for cDNA library preparation following manufacturer protocol. cDNA library was subject to quality control by Tapestation (HSD5000) and Qubit (High Sensitivity DNA) for mean size of ∼1500bp. Next, cDNA library was tagmented to prepare Illumina sequencing library (Illumina Nextera XT, UDI indexing). Quality control was performed with Qubit (High Sensitivity DNA) and Tapestation (HSD1000) for mean size of ∼300bp. Samples were loaded in equal molar with 3% spike-in PhiX DNA and sequenced on an Illumina NextSeq 1000 platform with paired-end 300bp cycle.

### Sequencing data preprocessing and quality control

Adapter sequences were trimmed using trimgalore (v0.6.7). Alignment was performed using STAR aligner^48^ (v2.7.0a) with Drosophila melanogaster dm6 genome and transcript annotation from the UCSC Genome Browser^49^. Parameter ‘--quantMode GeneCounts --outSAMtype BAM SortedByCoordinate’ was used. Read coverage and junction visualization of individual samples was performed with IGV^50^ (v2.16.1). Sashimi plots were generated by ggsashimi^51^ (v1.1.5). Gene-specific counts and alignment files were then used for downstream analysis.

### Sequencing data analysis

Differential gene expression analysis was performed with DESeq2^52^ (v1.46.0) following vignette standard procedures using gene-specific counts reported by the STAR aligner.

Differential alternative splicing analysis was performed with rMATS-Turbo^53^ (v4.3.0) using parameters “-t paired --allow-clipping --readLength 150 --variable-read-length --cstat 0.05” with dm6 genome (UCSC genome browser). Analysis results were further analyzed following published procedure of rMATS-Turbo. In brief, alternative splicing events were filtered to reject low-signal (minimum 10 reads) and extreme percent spliced-in (less than 5% or more than 95%) events. All criteria must be true for the mean values of both conditions sequenced. Filtered events are assigned up-/down-regulated or not-significant based on FDR < 0.1 and absolute percent spliced-in difference > 5% cutoffs. To quantify splicing of all introns in the genome, we used IRFinder-S^54^ (v2.0.1). Index was built with parameters “BuildRefProcess -j 100 -L 12000000000”. Alignment was quantified with “BAM -w 5”. “Spliced introns” are introns that are neither “LowCover” (less than 10 supporting reads) nor “LowSplicing” (less than 4 spliced junction reads) by IRFinder. Splicing ratio values from IRFinder-S was subsequently used for different downstream analysis described below.

### Classification of intron type in the Nascent-seq dataset reanalysis

Given a spliced intron, we classify it into four possible categories: “co-transcriptional splicing”, “post-transcriptional splicing -polyA spliced”, “post-transcriptional splicing - IntP-like”, and “other”. “co-transcriptional splicing” introns are those mean retention ratio is less than 50% in nascent RNA samples. “post-transcriptional splicing – polyA spliced” introns are those retained >80% in nascent samples, <40% in poly-A samples, and found to be statistically significant by IRFinder-S (padj < 0.01). “post-transcriptional splicing – IntP-like” introns are those retained >80% in nascent samples, between 40% to 70% in poly-A samples, and found to be statistically significant (padj < 0.01). If none of the above groups fit, an intron is assigned to the “other” group. Refer to Supplementary Table S1 for detailed results on intron splicing analysis.

### Intron splicing analysis of Prp3RNAi and control dataset

IRFinder was again used to analyze splicing of all introns. We identified 26003 introns in total (i.e., introns with at least 10 supporting reads). Next, introns are categorized into “constitutively spliced introns” (CS), “retained introns” (RI), or “others” (Other) with the control condition samples. CS introns have mean retention ratio <10% (found 22985, 88.4%). RI introns have mean retention ratio between 20%-80% (found 1617, 6.2%). If neither fit, introns put in Other category (5.4%).

A CS intron is “affected by Prp3-RNAi” if its mean retention ratio is >40% in Prp3-RNAi samples and IRFinder-S differential analysis p-value is less than 0.1. Out of 22985 CS introns, we found 38 (0.2%) to be more retained in Prp3RNAi. A RI intron is “affected by Prp3-RNAi” if its mean retention ratio is >40% in Prp3-RNAi samples and IRFinder-S differential analysis p-value is less than 0.1. Out of 1617 RI introns, we found 80 (4.9%) to be affected by Prp3-RNAi. 75 out of 80 (94%) affected introns are more retained in Prp3RNAi condition. Refer to Supplementary Table S4 for detailed results on intron splicing analysis.

### Characterization of intron properties

Splicing site strength scoring was computed with MaxEntScan^55^. Ratio of intron to mean exon length (RIME) and distance between inferred branchpoint position to the 3’ splice site was calculated with a custom R package following steps previously described by others^56^. A position weight matrix for consensus branchpoint was generated using the MEME suite^57^, and motif searching was performed with the Bioconductor universalmotif package. Precomputed evolutionary conservation phastCons scores were download from the UCSC Genome Browser (124 insect species multiway alignment) and mean scores along the full length of genomic regions were computed with a custom R package. Intron properties described is available in a R package ‘gsAnalysis’ (https://yeyuan98.r-universe.dev/gsAnalysis).

### Long-read sequencing

To extract RNA, we collected ∼150 fly heads by flash freezing in liquid nitrogen. Heads were homogenized in Trizol reagent. Total RNA was extracted, followed by DNase treatment (TURBO DNA-free Kit). Total RNA was measured by Nanodrop and Qubit. polyA+ enrichment was performed (NEBNext High Input PolyA mRNA Isolation Module) and enriched RNA was measured by Nanodrop and Qubit. This procedure usually yielded ∼200ng polyA-enriched mRNA (∼2% of total RNA before enrichment). To prepare Nanopore sequencing library on a DNA flow cell, we used Oxford Nanopore cDNA-PCR Kit v14. Briefly, 10ng enriched RNA is reversed-transcribed by template switching using mixture of oligo-dT and target-specific (tim and Sxl) primers. Product is subsequently amplified with manufacturer cDNA Primers to enrich full-length transcripts and subsequent adapter addition. The prepared library is subsequently loaded on a MinION Flow Cell (R10.4.1) and sequenced with the MinKNOW controller software for ∼8hr. To analyze sequencing data, we performed base calling and alignment with Nanopore Dorado with HAC basecalling model and UCSC dm6 genome. Alignment is filtered with using custom R/Bioconductor script to look for reads aligning to the desired region with sufficient read length. Filtered reads are visualized with the IGV genome browser. Filtering criteria to select long mapped reads specific to timeless is as follows: 1) one end is in the last exon of timeless “long isoforms”, 2) reference map length is >8kb, 3) maximum intron length is <4kb, 4) maximum indel length is 100bp.

### Sequence conservation analysis

While phastCons scoring reflects per-base sequence conservation (see Methods ‘Characterization of intron properties’), to further evaluate conservation of intron P in genus Drosophila, we performed BLAST search of intron P (database: Refseq Reference genomes, organism: Drosophila 7215, blastn algorithm). The resulting multiple sequence alignment was exported to as an ‘aln’ file and visualized using the JalView^58^ program with consensus and occupancy visualization tracks.

### Locomotor activity analysis

Individual adult male flies (3-5 days old) were placed in glass capillary tubes containing 2% agar and 4% sucrose food and loaded into TriKinetics DAM2 Drosophila Activity Monitors (Waltham, MA, USA) for locomotor activity recordings. Flies were entrained to Light-Dark (LD) cycles with lights on for 12 hours and off for 12 hours for 5 days, followed by complete darkness (DD) for 7 days. DAM monitors record infrared beam breaks when the flies cross the middle of the glass tubes as 30-minute binned reading. Averaged population activity profiles (actograms) under Light-Dark cycles and constant conditions were generated using R Rethomics package. Briefly, activity levels were normalized for each fly, such that the average number of beam crossings in each day (48 bins) is equal to 1. Next, the population average of normalized activity was determined and the results plotted as actogram figures. We used activity counts of individual flies under DD conditions to determine the free-running period and amplitude of the circadian clock by a chi-square periodogram analysis with a confidence level of 0.001 and period cutoffs 18-30 hours using the ClockLab software. The “Power” and “Significance” values generated from the chi-square analysis were used to calculate “Rhythmic Power” to reflect oscillation strength. Custom functions for actogram plotting and batch periodogram summary is available in a R package ‘ijAnalysis’ (https://yeyuan98.r-universe.dev/ijAnalysis).

### Estimating splicing rate of intron P

To estimate the effect of intron P on reporter RNA processing (Figure 3l-m), we adopted an existing two-state kinetics model^35^. In brief, transcription (T) yields pre-mRNA (*P*) in the nucleus, which is processed (spliced and exported, with a rate constant k_ps_) into mature mRNA (M) in the cytoplasm. Both pre-mRNA (P) and mature mRNA (M) are subject to degradation (rate constants k_p_ and k_m_, respectively). The following equations describe the time variation of P and M (assuming first-order reactions):

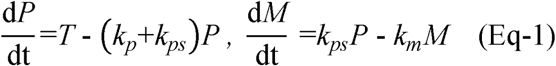

In the reporter assay (Figure 3), *Clk-GAL4* driver is used for expression of reporter minigenes (Exon and IntP) positioned at the attP2 genomic site. Hence, transcription (T) is identical for both reporters. Further, cytoplasmic degradation (k_m_) is identical, as the mature transcript is always mCherry-NLS-Exon-tetR. Finally, the system is at steady state, given the constant overexpression of the reporter and lack of feedback mechanisms.

With the above assumptions, nuclear enrichment (named as ‘NE’, NE=P/(P+M)) depends solely on *k_m_* and *k_ps_*, while the total reporter level (named as ‘TL’, TL=P+M) depends on all kinetic parameters (*k_p_, k_m_* and *k_ps_*) as shown below. Both NE and TL can be measured experimentally. Under stead state conditions:

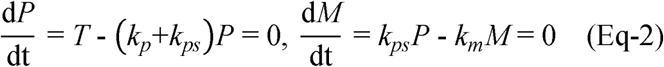

which yields:

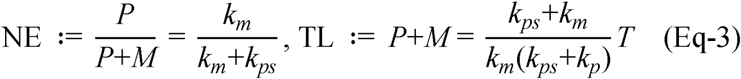

Compared to TL which relies on complex parameters, NE provides a direct readout of k_ps_/k_m_:

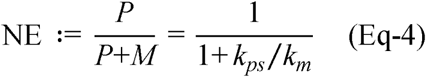

For instance, when NE is measured to be 25%, we have k_ps_/k_m_*=*3. Considering that *k_m_* is a constant and *k_ps_* depends on the reporter construct (i.e., Exon and IntP constructs have k_ps,Exon_ or k_ps,E_ and k_ps,IntP_ or k_ps,I_), by comparing NE values from different reporters, we can get the ratio of *k_ps_* between different reporter constructs.

Estimates provided in Fig. 3m are based on ZT4 data of Fig. 3c. Considering first and third quantiles of the measured NE, Exon reporter shows NE of 22.0-28.7%, and IntP reporter shows NE of 70.8∼78.0%. This yields k_ps,Exon_*/k_m_*∼ 2.5-3.6 and k_ps,IntP_*/k_m_* ∼ 0.3-0.4. Hence, we get k_ps,Exon_/k_ps,IntP_ ∼ 6-13, suggesting that intron P is inefficiently spliced.

### Numerical simulation of the circadian clock

We implemented a previously established 10-state circadian clock model^24, 59^ in R using the Odin package for efficient numerical simulation. In brief, the clock model consists of 10 ordinary differential equations (ODE) describing a negative feedback loop created by PER-TIM gene transcription, mRNA translation, protein multistep phosphorylation (un-, under-, and hyper-phosphorylated), PER-TIM complex formation (only when PER and TIM proteins are hyper-phosphorylated) and nuclear entry. These 10 ODEs were solved using the deSolve R package^60^ with the variable-step lsoda solver under default settings, which confirmed numerical convergence of all simulation runs. The model emulates light entrainment of the clock via light-dependent degradation rate of TIM protein (doubled rate when lights are turned on). Therefore, to entrain the model to a fixed light schedule and synchronize different simulations, we first subject the model to 20 days of LD12/12 cycle before releasing in constant DD conditions. Numerical integration was performed for a total of 50 days with 1-hour output time intervals, and *tim* mRNA data from the last 10 days of the simulation were subject to JTK Cycle^61^ rhythmicity analysis to get period and amplitude of *tim* mRNA oscillation. To understand whether the circadian clock system behaves as a limit cycle oscillator in normal and mutant conditions, we plot trajectory of the system with *tim* RNA M_T and nuclear PER-TIM complex C_N as X and Y axis (i.e., phase portrait plot). Normally, the molecular clock behaves as a limit cycle oscillator, which explains its fixed-period and self-sustained oscillation properties^62^. Under the limit cycle condition, the system should follow a determined trajectory, forming a “circle” in the phase portrait plot. The numerical simulation workflow is available as a R package ‘clockSim’ (https://cran.r-project.org/web/packages/clockSim).

## Acknowledgments

We thank members of the Yadlapalli laboratory, Pramod Reddy, Edgar Meyhofer, and Nils Walter for helpful discussions on the manuscript. We thank Amirah Nieves for help with some experiments. We thank Joanna Chiu for sharing the anti-TIM antibody. We thank Catherine Hsieh and Sami Barmada for initial help with U2OS cell culture experiments. We thank Bowen Yan, Hao Wu, and Zongyuan Liu for discussions on numerical simulation. We acknowledge support from the University of Michigan Biomedical Research Core Facilities (Flow Cytometry Core and Advanced Genomics Core). We thank the Bloomington Drosophila Stock Center (funded by NIH grant P40 OD018537) for providing fly strains.

## Author contributions

Conceptualization: SY

Methodology: YY, AL, SY

Investigation: YY, AL, RDG, HL, YX, SS, IH, SY

Visualization: YY, AL, RDG

Funding acquisition: SY

Project administration: SY

Supervision: SY

Writing – original draft: YY, SY

Writing – review & editing: YY, AL, SY

## Data and materials availability

All data are available in the main text or the supplementary materials. Sequencing alignment can be found at the Gene Expression Omnibus (accession: GSE293258). Raw data is available at the Sequence Read Archive (accession: PRJNA1243426).

## Supplementary Materials

Extended Data Figs. 1 to 10

Supplementary Tables S1 to S10

---

**Extended Data Figure 1.**
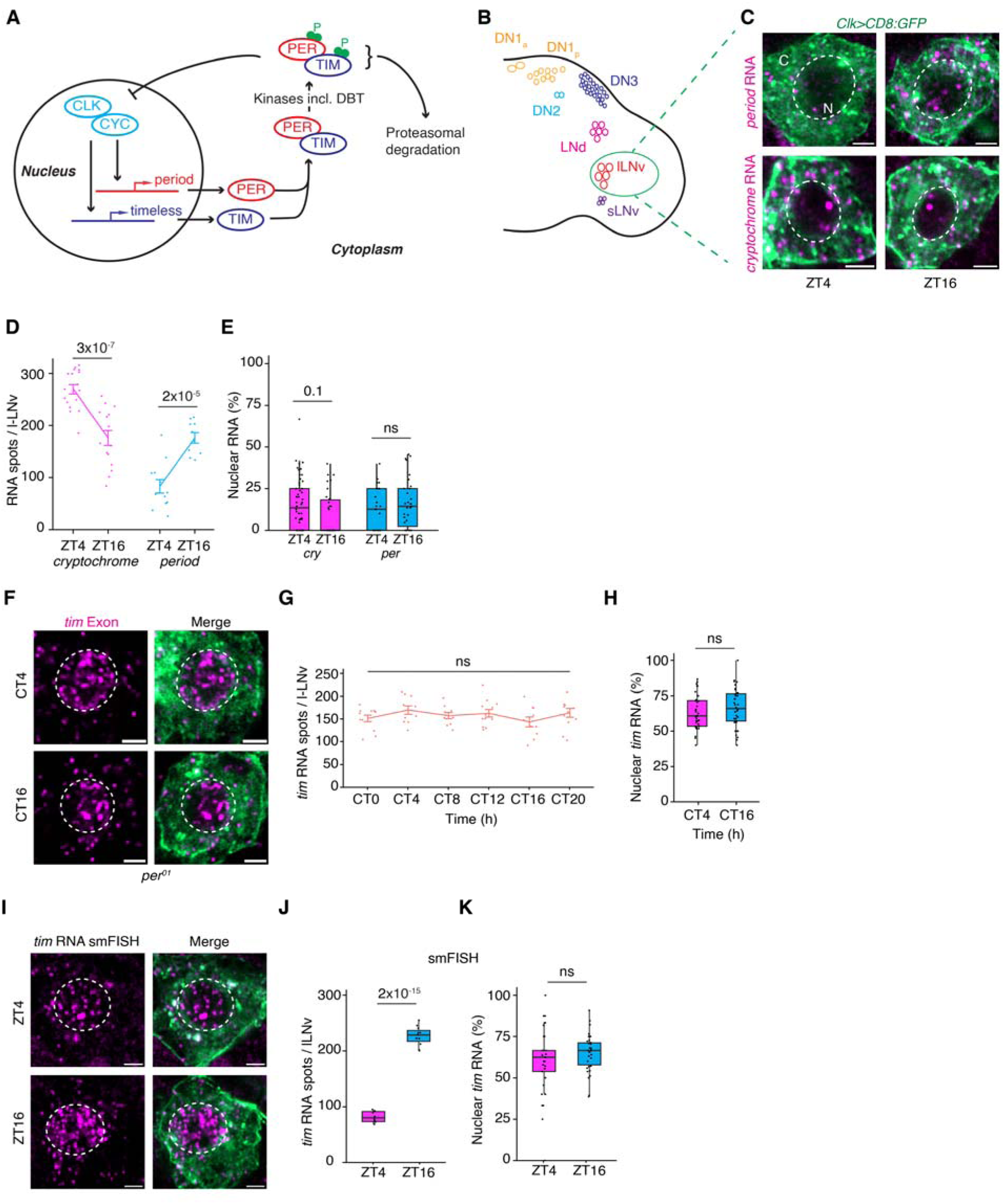
*Timeless* mRNAs are localized to both the nucleus and cytoplasm of clock neurons. (**A**) Schematic of the *Drosophila* molecular clock. Briefly, In *Drosophila*, the CLOCK–CYCLE complex activates the transcription of many genes, including the core clock genes *period* (*per*) and *timeless* (*tim*), which encode repressors. The PER and TIM proteins gradually accumulate, and, after a time delay, translocate to the nucleus and inhibit their own transcription as well as that of other clock-controlled genes. Upon the degradation of PER and TIM proteins, the activation phase resumes, completing a feedback loop that takes ∼24 hours. (**B**) Schematic of clock neuron groups in a fruit fly hemi-brain. (**C**) Representative images of *cryptochrome* (*cry*) and *period* (per) RNA in l-LNvs during the repression (ZT4) and the activation (ZT16) phases. l-LNv cytoplasm is shown in green, RNA spots are shown in magenta, and nuclei are outlined in white. (**D, E**) Quantification of total *cry* and *per* RNA spots (**D**) and percentage of nuclear RNA (**E**) in l-LNvs. (**F**) Representative images of *tim* RNA in l-LNvs from *per^01^* null mutant flies. (**G**) Quantification of *tim* RNA spot count over the first day of darkness (DD1). In *per^01^* mutant flies, *tim* RNA count remains high (near peak value of wildtype) throughout the circadian cycle. (**H**) Quantification of percentage of nuclear *tim* RNA in l-LNVs from *per^01^* flies. (**I**) Representative images of single molecule RNA-FISH (smFISH) of *tim* RNA in l-LNvs during repression (ZT4) and activation (ZT16) phases. (**J**) Quantification of *tim* RNA spot count (smFISH) per l-LNv. (**K**) Quantification of percentage of nucelar *tim* RNA (smFISH) in l-LNvs. In **D**, and **G,** data are presented as line plots showing the mean ± standard error of the mean (SEM). Statistical analysis was performed using a two-sided Student’s t-test. In **E**, **H**, **J**, and **K**, data are shown as box plots displaying the median and quantiles. Statistical analysis was performed using two-sided Mann-Whitney U tests. In all plots, each dot represents data from l-LNvs of a single *Drosophila* hemi-brain. Refer to Supplementary Table S10 for all statistical comparisons. l-LNv nuclei are circled in white. Scale bars-2µm.

**Extended Data Figure 2.**
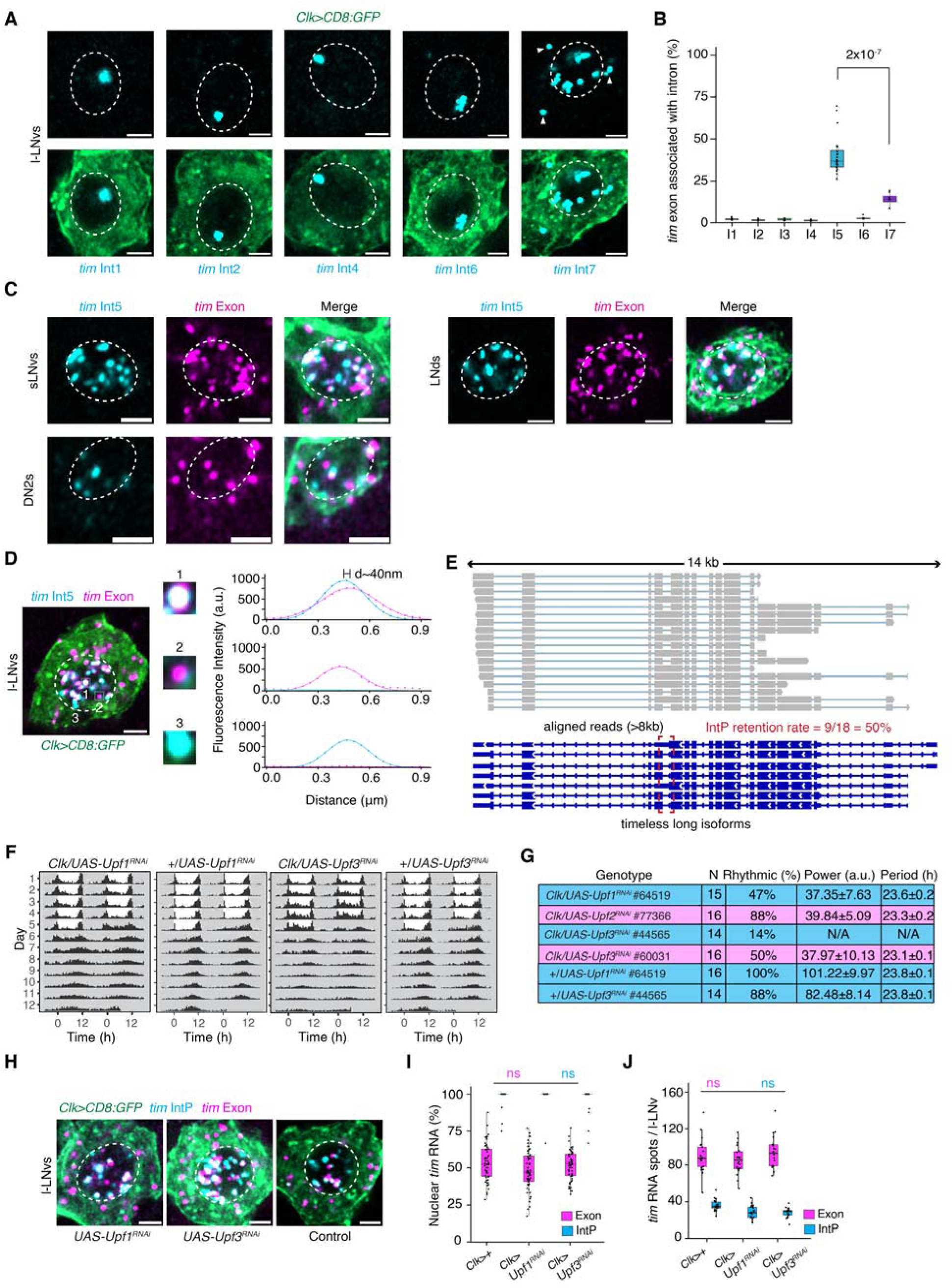
*tim* mRNAs are spliced post-transcriptionally. (**A**) Representative images of FISH signal (cyan) for various *tim* introns in l-LNvs. The FISH signal for most introns (except intron 5 and 7) appears as a single nuclear spot, consistent with co-transcriptional splicing. Intron 7, which has been previously shown to be alternative spliced^21^; shows both cytoplasmic and nuclear signals. (**B**) Percentage of *tim* exon spots (both nuclear and cytoplasmic) colocalizing with intron spots in l-LNvs. (**C**) Representative images of *tim* exon and intron 5 in different clock neuron subgroups. (**D**) Representative image showing FISH signals for the *tim* exon and intron 5. Line intensity traces on the right illustrate examples of a colocalized exon-intron 5 spot (top), an exon-only spot (middle), and an intron 5-only spot (bottom). (**E**) Long-read sequencing reads aligned to the *tim* gene locus (see Methods). Shown here are reads that start at the 3′ end of the long *tim* isoforms and span more than 8 kb of the genome. Of the 18 such reads, 9 retain intron 5, indicating a retention rate of ∼50%. (**F**) Averaged population locomotor-activity profiles of flies (*Clk>Upf1-RNAi*, *Clk>Upf3-RNAi*, and the corresponding controls) in LD and DD conditions, with rest-activity patterns shown for two consecutive days in each row. (**G**) Periodogram analysis of flies in constant conditions. (**H-J**) Representative images of *tim* exon and IntP FISH signals (**H**), quantification of percentage of nuclear *tim* exon and intron P signals (**J**), quantification of *tim* exon and intron P spot count (**J**) in l-LNvs under Upf1 and Upf3 knockdown and control conditions. IntP spots are exclusively localized in the nucleus. There is no significant difference observed in any of the analyses across the three conditions. ns, not significant. In **B**, **I**, and **J**, data are shown as box plots displaying the median and quantiles. Statistical analysis was performed using two-sided Mann-Whitney U tests. In all plots, each dot represents data from l-LNvs of a single *Drosophila* hemi-brain. Refer to Supplementary Table S10 for all statistical comparisons. l-LNv nuclei are circled in white. Scale bars-2µm.

**Extended Data Figure 3.**
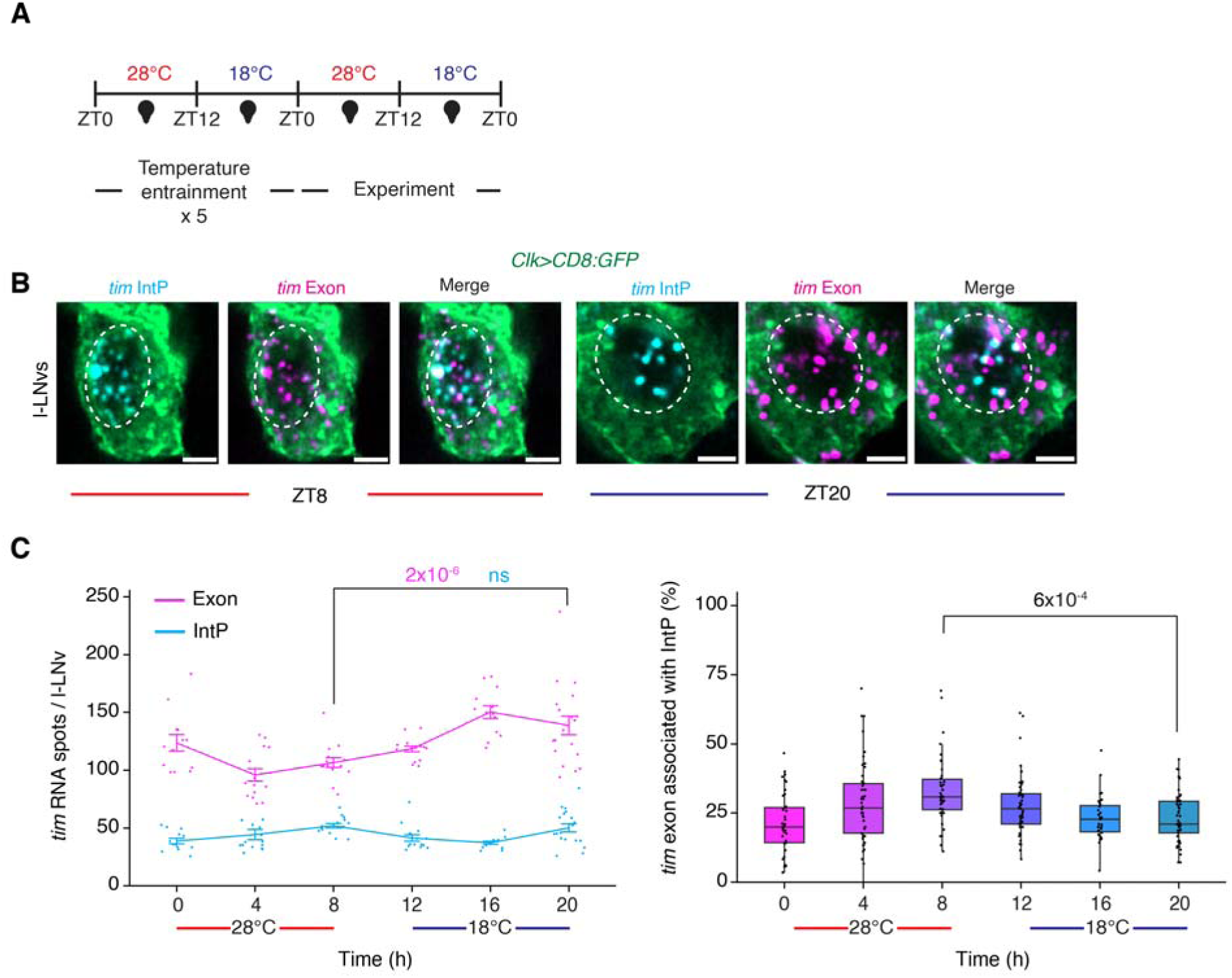
Intron P splicing is slightly more efficient at lower temperatures. (**A**) Schematic of temperature cycle entrainment. Flies were kept under constant darkness and subject to cycling temperature conditions (28 °C/18 °C). ZT0 denotes the start of the warm phase (28 °C), ZT12 denotes the start of the cold phase (18 °C). (**B**) Representative image of *tim* exon and intron P FISH signals in l-LNvs at different time points over the temperature cycle. ZT8 corresponds to a timepoint in the warm phase, while ZT20 corresponds to a timepoint during the cold phase. (**C**) Quantification of the total number of *tim* exon spots (pink trace) and intron P spots (blue trace) per l-LNv at different times over the temperature cycle is shown in the panel on the left. P-values for RNA levels at ZT8 vs. ZT20 are shown in pink for the exon and in blue for intron P. ns; not significant. Data are presented as line plots showing the mean ± standard error of the mean (SEM). Statistical analysis was performed using a two-sided Student’s t-test. Quantification of the percentage of nuclear *tim* exon signal per l-LNv across the temperature cycle is shown in the panel on the right. Data are shown as box plots displaying the median and quantiles. Statistical analysis was performed using two-sided Mann-Whitney U tests. In all plots, each dot represents data from l-LNvs of a single *Drosophila* hemi-brain. Refer to Supplementary Table S10 for all statistical comparisons. l-LNv nuclei are circled in white. Scale bars-2µm.

**Extended Data Figure 4.**
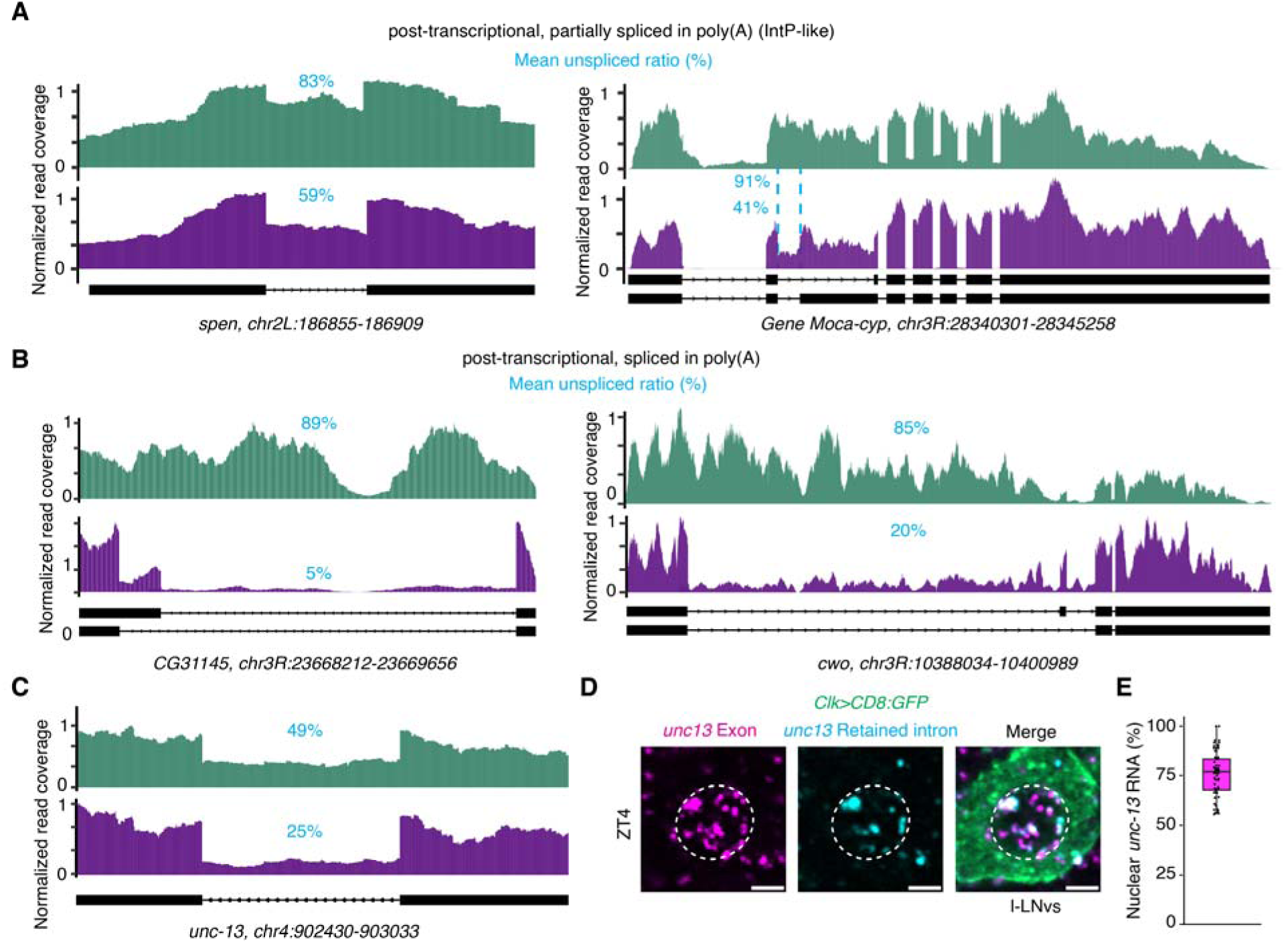
Analysis of Nascent-sequencing data from Drosophila brains. (**A**) Normalized read coverage of Nascent-seq samples (top) and poly(A)-seq samples (bottom) for representative loci that are characterized as “post-transcriptional splicing, IntP-like”, which are introns that are retained to high levels in both nascent and poly(A) transcripts (Supplementary Table S1). (**B**) Normalized read coverage for representative loci characterized as “post-transcriptional splicing, spliced in poly(A)”, which are introns that are present in nascent RNA but spliced out in poly(A) transcripts. (**C**) Normalized read coverage for an intron of *unc-13* that is retained in both nascent RNA and poly(A) RNA. Splicing status of the respective introns, quantified by mean unspliced read ratio (averaged over three replicates) is shown as percentages in blue. (**D**) Representative image of *unc-13* exon and intron (which is retained in the nucleus) spots in l-LNVs. *unc-13* is not exclusive to clock neurons and therefore signals from neighboring cells are visible. **e)** Quantification of the percentage of nuclear *unc-13* exon spots in l-LNvs. Data are shown as box plots displaying the median and quantiles, each dot represents data from l-LNvs of a single *Drosophila*hemi-brain. Scale bars-2µm.

**Extended Data Figure 5.**
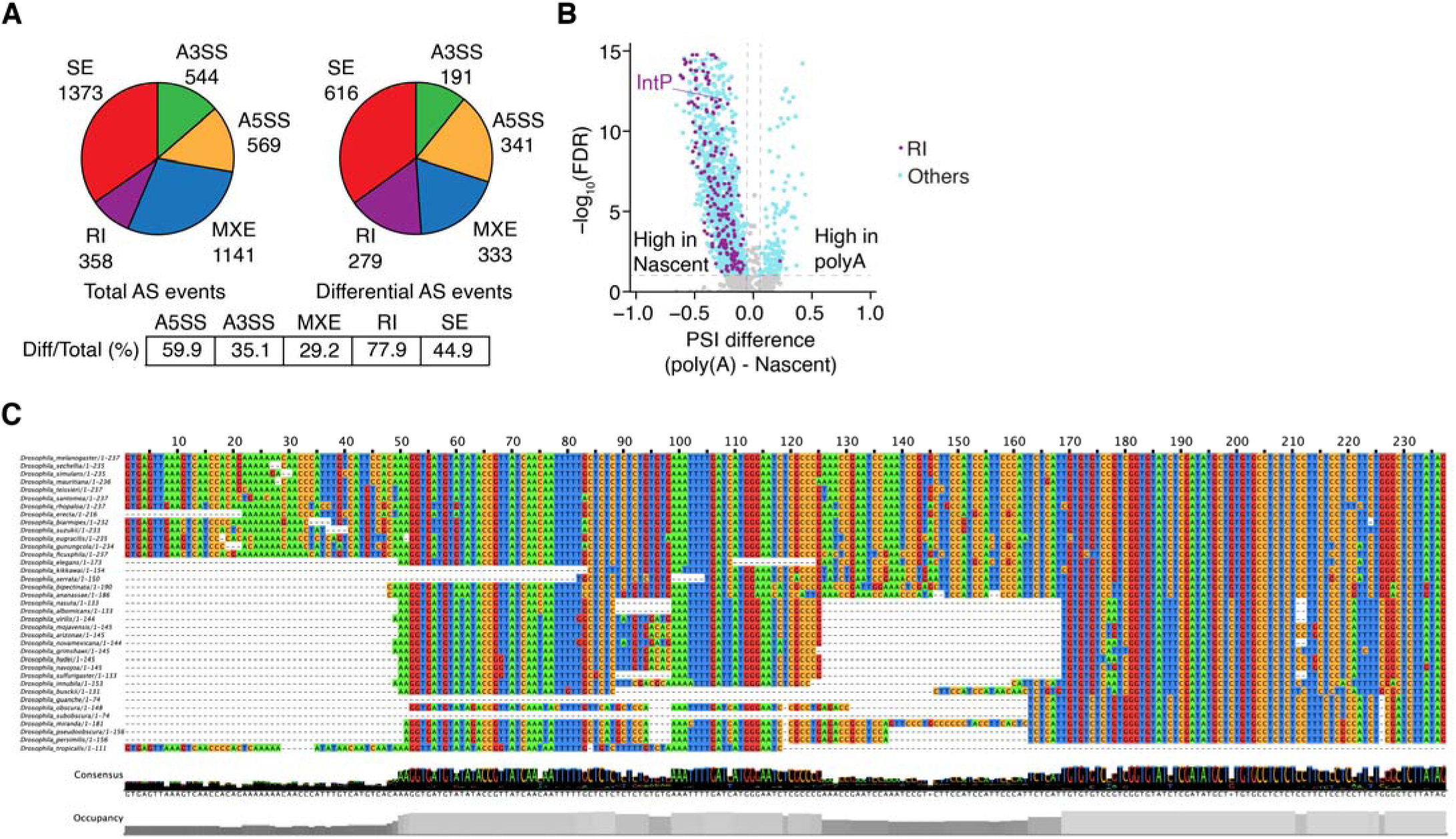
Analysis of alternatively spliced introns using the Nascent-sequencing dataset. (**A**) Pie chart showing the differential analysis of alternative splicing (AS) events between Nascent-seq and poly(A) RNA samples (Supplementary Table S2). Categories include A5SS: Alternative 5’ Splice Site, A3SS: Alternative 3’ Splice Site, MXE: Mutually Exclusive Exon, RI: Retained Intron, SE: Skipped Exon. (**B**) Volcano plot of differential AS events, with significant events shown in cyan and significant retained intron events in purple. (**C**) Multiple sequence alignment of *tim* intron P across different *Drosophila* species. Intron P is conserved in *timeless* gene homologs in most *Drosophila* species.

**Extended Data Figure 6.**
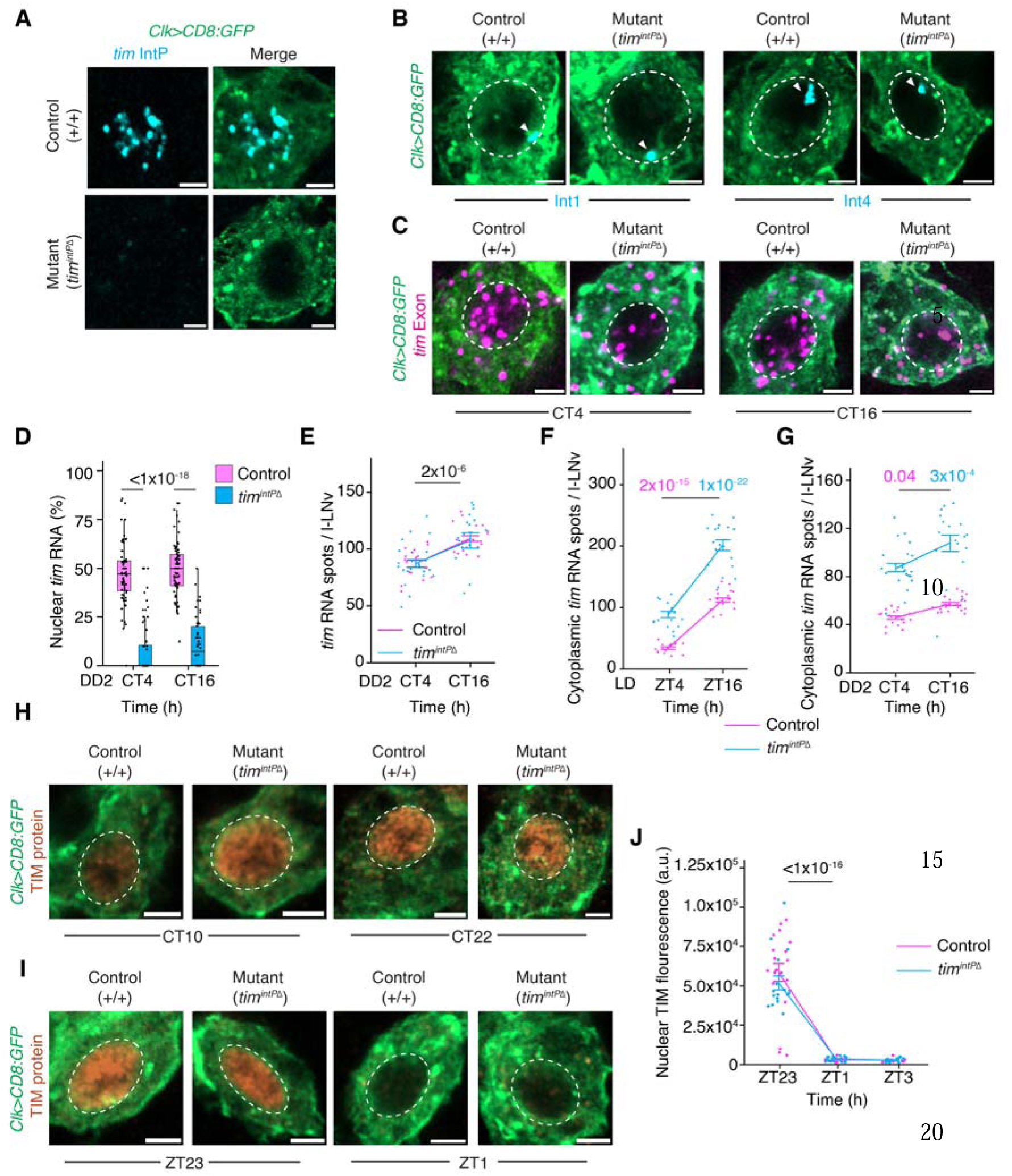
**Analysis of the *tim^IntP^***^Δ^ **CRISPR fly line.** (**A**) Representative images of intron P (cyan) in l-LNvs (cytoplasm labeled in green) from wild-type control and *tim^IntP^*^Δ^ flies. No signal is detected in *tim^IntP^*^Δ^, indicated successful CRISPR-mediated knockout of intron P. (**B**) Representative images of introns 1 and 4 in l-LNvs from control and *tim^IntP^*^Δ^ flies. (**C-E**) Representative images of *tim* RNA (**C**) quantification of nuclear tim RNA (**D**), and total number of tim mRNA spots (**E**) in l-LNVs from control and *tim^IntP^*^Δ^ flies at two timepoints during the second day of constant darkness (DD2). (**F, G**) Quantification of cytoplasmic *tim* RNA spot count in l-LNvs under LD (**F**) and DD2 (**G**) condition. P-values for RNA levels at ZT4 versus ZT16 are shown in pink for control and in blue for *tim^IntP^*^Δ^ flies. (**H**) Representative images of TIM protein in l-LNvs from control and *tim^IntP^*^Δ^ flies during repression (CT22) and activation (CT10) phases. (**I**) Representative images of TIM protein immunofluorescence in l-LNvs from control and *tim^IntP^*^Δ^ flies, taken one hour before (ZT23) and one hour after (ZT1) lights are turned on. (**J**) Quantification of TIM protein fluorescence intensity in l-LNvs. Light-dependent degradation of TIM protein is not affected in *tim^IntP^*^Δ^ flies. In **E, F**, **G**, and **J**, data are presented as line plots showing the mean ± standard error of the mean (SEM). Statistical analysis was performed using a two-sided Student’s t-test. In **D**, data are shown as box plots displaying the median and quantiles. Statistical analysis was performed using two-sided Mann-Whitney U tests. ns; not significant. In all plots, each dot represents data from l-LNvs of a single *Drosophila* hemi-brain. Refer to Supplementary Table S10 for all statistical comparisons. l-LNv nuclei are circled in white. Scale bars-2µm.

**Extended Data Figure 7.**
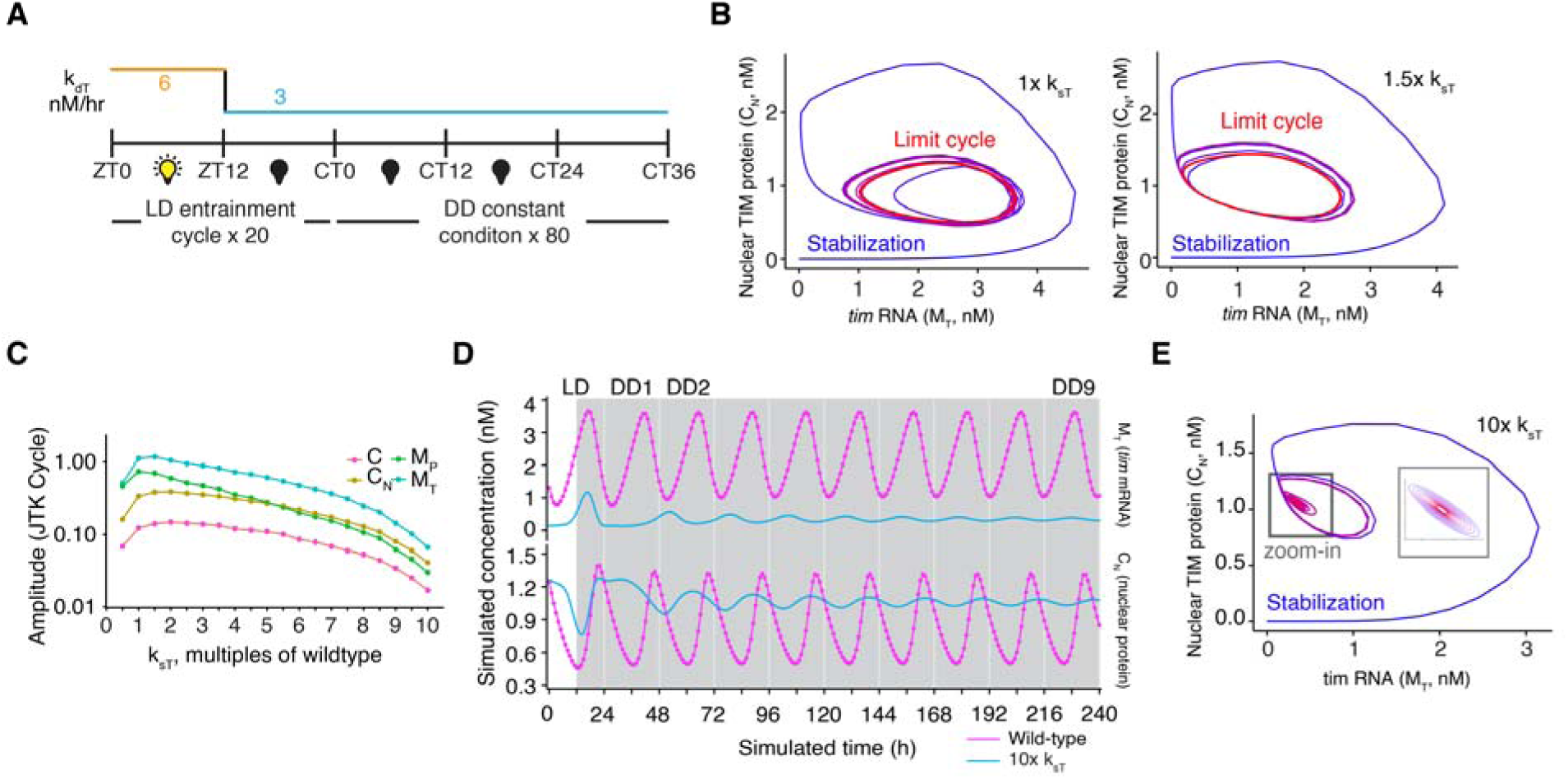
Numerical simulation of the molecular clock in intron P deletion condition. (**A**) Schematic of the numerical simulation setup. The model simulates a light–dark cycle by time-dependent modulation of the TIM protein degradation rate. It first runs for 20 days under a 12-hour light : 12-hour dark cycle, followed by release into constant conditions. To quantify the period and amplitude of the clock, JTK Cycle analysis was performed on the final 10 days of the constant condition phase. The total simulation duration was 50 days. (**B**) Limit cycle oscillations were observed using the default parameters,1.0x, from the original model (left) and when the translation rate of *tim* mRNA was increased to 1.5x (right). ‘k_sT_’ refers to the translation rate of *tim* mRNA. (**C**) Periodogram analysis showing oscillation amplitude at different TIM protein translation rates. The *tim* mRNA (M_T) state was used for this analysis; other molecular states are synchronous but exhibit different phase lags. Increasing the TIM protein translation rate leads to damped oscillations of *per/tim* mRNA and PER-TIM protein complexes. ‘C’: Cytoplasmic PER-TIM complex, ‘C_N_’: Nuclear PER-TIM complex, ‘M_p_’: *per* mRNA, and ‘M_t_’: *tim* mRNA. (**D**) Time series of the clock model when the *tim* mRNA translation rate is increased 10-fold. (**E)** Under the 10xk_sT_ condition, the model converges to a stable point after releasing into constant conditions, indicating a disrupted clock (*i.e.*, loss of limit cycle oscillation). See Methods on additional details about this simulation.

**Extended Data Figure 8.**
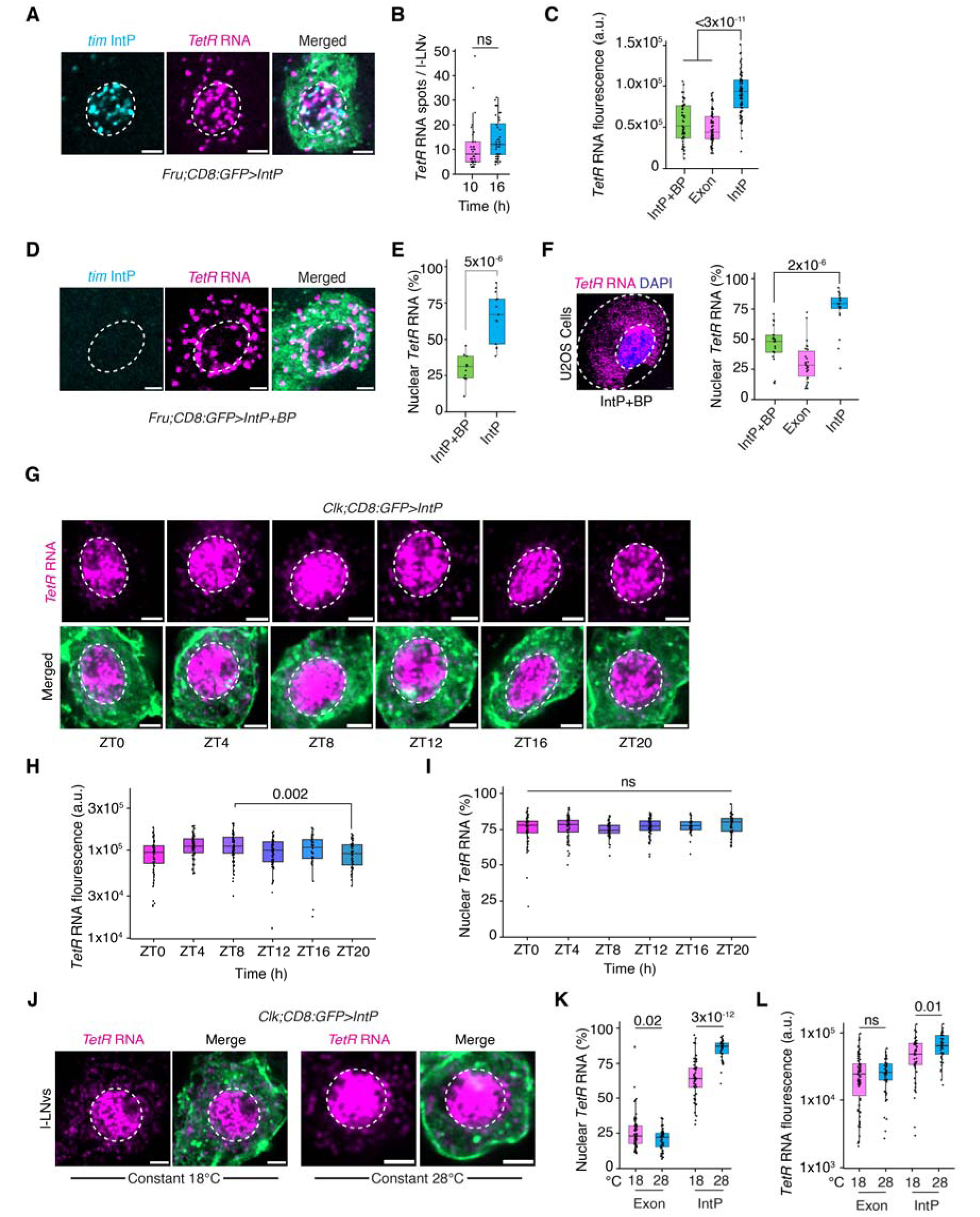
Intron P is sufficient for nuclear enrichment of reporter mRNAs in *Drosophila* neurons and human U2OS cells. (**A**) Representative image of *TetR* RNA in *Drosophila* courtship neurons (*Fru-GAL4*-positive) from IntP flies. (**B**) Quantification of total *TetR* RNA spots in l-LNvs from IntP flies at 10 and 16 hours after flies were shifted to 29 °C to induce *tetR* RNA expression. (**C**) Quantification of total *TetR* levels in l-LNvs from Exon (control), IntP, and IntP+BP flies. (**D**) Representative image of *TetR* RNA and intron P in *Drosophila* courtship neurons from IntP+BP flies. (**E**) Quantification of the percentage of nuclear *TetR* RNA in *Drosophila* courtship neurons from IntP and IntP+BP flies. (**F**) Representative image of *TetR* RNA in human U2OS cells transfected with IntP+BP construct (left panel). Quantification of the percentage of nuclear *TetR* RNA in U2OS cells transfected with Exon, IntP, and IntP+BP constructs (right panel). (**G**) Representative images of *TetR* RNA in l-LNVs from IntP flies over the light-dark cycle. (**H, I**) Quantification of total *TetR* RNA **(H)** and percentage of nuclear *TetR* RNA **(I)** in l-LNVs over the light-dark cycle. (**J**) Representative images of *TetR* RNA in l-LNVs from IntP flies at constant low (18 °C) or high (28 °C) temperatures. (**K, L**) Quantification of percentage of nuclear *TetR* RNA **(K)** and total *TetR* RNA **(L)** in l-LNvs from IntP flies at 18 °C and 28 °C. *TetR* RNA shows higher cytoplasmic localization at the lower temperature (18 °C) compared to the higher temperature (28 °C). In **B**, **C**, **E**, **F**, **H**, **I**, **K**, and **L**, data are shown as box plots displaying the median and quantiles. Statistical analysis was performed using two-sided Mann-Whitney U tests. ns; not significant. In all plots, each dot represents data from l-LNvs of a single *Drosophila* hemi-brain. Refer to Supplementary Table S10 for all statistical comparisons. Nuclei are circled in white. Scale bars-2µm.

**Extended Data Figure 9.**
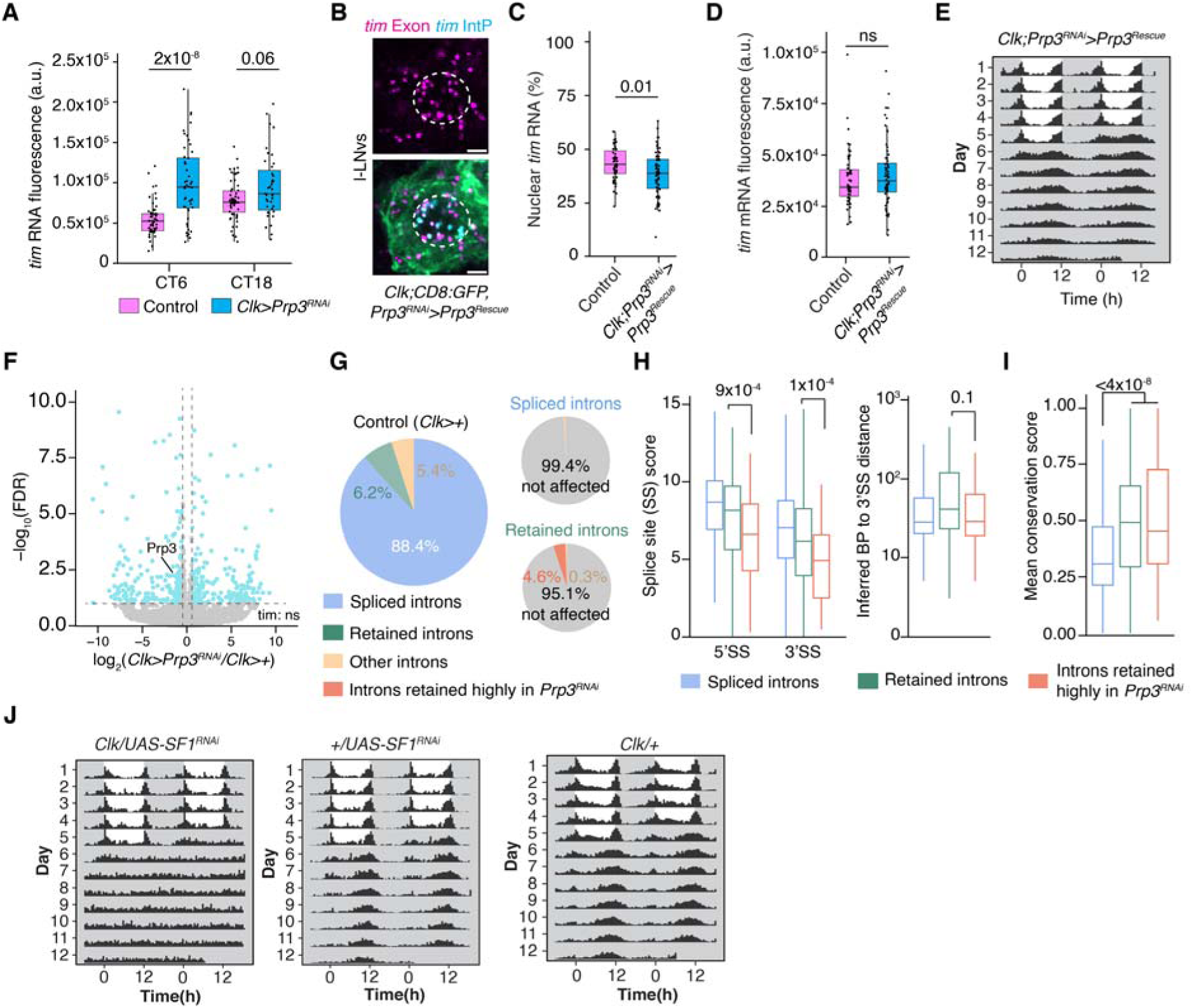
**Splicing of intron P is specifically by affected by knockdown of spliceosome factors, including Prp3, SF1 and Prp19**. (A) Quantification of total *tim* RNA fluorescence in l-LNvs from control and *Clk>Prp3-RNAi* flies. Knockdown of Prp3 leads to increased *tim* RNA levels. (B) Representative images of *tim* exon and intron P in Prp3-RNAi with overexpression rescue with Prp3-V5. (C) Quantification of percentage of nuclear *tim* RNAs in overexpression rescue flies. (D) Quantification of *tim* RNA in overexpression rescue flies. Overexpression of Prp3 in knockdown background rescues both behavioral and molecular clock defects of Prp3 knockdown. (E) Behavior actogram of the *Prp3-RNAi/Prp3-V5* knockout rescue experiment. (F) Volcano plot of differential gene expression analysis. Prp3 is significantly down-regulated in Prp3-RNAi knockdown to ∼50% of wildtype levels. Overall mostly balanced up-/down-regulation is observed without clear gene ontology enrichment indicating diverse targets of Prp3 (Supplementary Table S6). (G) Pie chart of differential intron splicing analysis. In wildtype conditions, most introns are not retained (‘Group 1-Spliced Intron’) while 6.2% is retained (‘Group 2-Retained Intron’). Prp3 knockdown has little effect on Group 1 introns as >99% remains spliced. Prp3 knockdown affects a small portion (4.9%) of Group 2 introns, the majority of which (4.6%, designated as Group 3 Introns) show higher unspliced ratio upon knockdown, indicating that these introns require higher levels of Prp3 for splicing (Supplementary Table S4). (H) Quantification of intron splice site strength (MaxEnt score) and distance from inferred branch point to 3’ splice site of Group 1-3 introns. (I) Quantification of mean conservation score (PhastCons). (J) Behavior actogram of parental control and SF1-RNAi knockdown conditions. Statistical tests are two-sided Mann-Whitney U test. Refer to Supplementary Table S10 for statistics. Nucleus is circled in white. Scale bars-2µm.

**Extended Data Figure 10.**
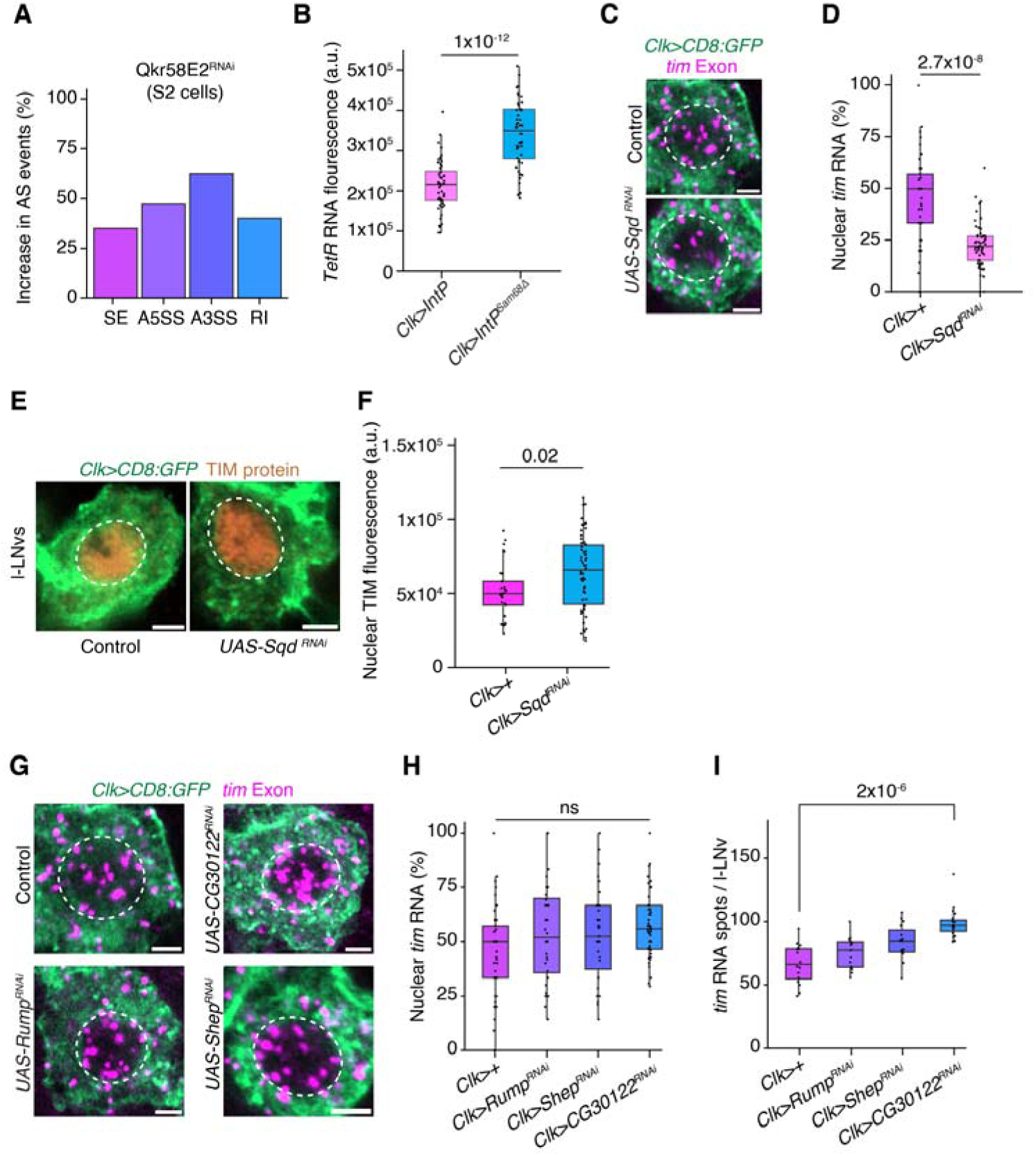
A genetic screen of splicing regulator RBPs uncovers activators and repressors of intron P splicing. (**A**) Knockdown of Qkr58E-2 in *Drosophila* S2 cells affects all major AS types and can act as both an activator and repressor. Categories include A5SS: Alternative 5’ Splice Site, A3SS: Alternative 3’ Splice Site, MXE: Mutually Exclusive Exon, RI: Retained Intron, SE: Skipped Exon. (**B**) Quantification of *TetR* RNA intensity in l-LNvs from the control (*IntP*) and motif-deleted (*IntP^Sam68^*^Δ^) reporter fly lines. (**C**) Representative images of *tim* RNA in l-LNvs from control and *Clk>Sqd-RNAi* flies. (**D**) Quantification of percentage of nuclear *tim* RNA in l-LNvs from control and *Clk>Sqd-RNAi* flies. (**E**) Representative images of TIM protein in l-LNvs from control and *Clk>Sqd-RNAi* flies. (**F**) Quantification of TIM protein fluorescence intensity in l-LNvs from control and *Clk>Sqd-RNAi* flies. (**G**) Representative images of *tim* RNA in l-LNv neurons at repression phase (CT0) in CG30122/Rump/Shep knockdown and control conditions. (**H, I**) Quantification of percentage of nuclear *tim* RNA (**H**) and total number of *tim* transcripts **(I**). In **B**, **D**, **F**, **H**, and **I**, data are shown as box plots displaying the median and quantiles. Statistical analysis was performed using two-sided Mann-Whitney U tests. ns; not significant. In all plots, each dot represents data from l-LNvs of a single *Drosophila* hemi-brain. Refer to Supplementary Table S10 for all statistical comparisons. Nuclei are circled in white. Scale bars-2µm.

**Table S1. Analysis of intron splicing from Nascent-seq and poly(A)-seq datasets**

**Table S2. Analysis of alternative splicing from Nascent-seq and poly(A)-seq datasets**

**Table S3. Behavioral screening of spliceosomal factors**

**Table S4. Intron splicing analysis from Nascent RNA-seq data**

**Table S5. Alternative splicing analysis from Nascent RNA-seq data**

**Table S6. Differential gene expression analysis**

**Table S7. Behavioral screening of splicing regulatory RBPs**

**Table S8. Behavioral results of control lines**

**Table S9. Plasmid and gRNA sequences**

**Table S10. Statistical analysis**

